# Mesolimbic dopamine signaling mediates increased hedonic feeding and food seeking in lactating mice

**DOI:** 10.64898/2026.03.27.714897

**Authors:** Tanya Pattnaik, Benjamin Wang, Emma Borrowman, Vraj Patel, Qingying Zheng, Lara Villano, Patrick Sweeney

## Abstract

Lactation dramatically increases energy intake to support milk production and care for the offspring. However, the behavioral and neural circuit mechanisms driving heightened feeding during lactation remain unclear. Here, we reveal that lactation increases food-seeking behavior and enhances palatable food intake in mice. Fiber photometry recordings demonstrate increased dopamine release in the nucleus accumbens in lactating animals during feeding tasks. This elevated dopamine signaling is ultimately required for promoting both food seeking and palatable food intake during lactation as pharmacological inhibition of dopamine receptors or chemogenetic inhibition of VTA dopamine neurons both reduce food seeking and palatable food intake in lactating mice to non-lactating levels. Further, selective inhibition of dopamine receptors in the nucleus accumbens produces similar results. Together, these findings provide a circuit basis mediating elevated food seeking and palatable food intake during lactation, providing novel insights into the regulation of maternal energy balance and feeding behavior.

**Highlights:** Lactation increases food seeking and hedonic feeding in mice

Mesolimbic dopamine levels are enhanced during feeding in lactating mice

VTA dopamine neurons mediate increased palatable food intake and food seeking during lactation

VTA-NAc dopamine transmission mediates increased food seeking and palatable food intake during lactation

## Introduction

The energetic demands of milk production and caring for young present an enormous metabolic challenge in mammals which is met by drastically increasing food intake^1^. For example, food intake increases 200-400 percent in rodents during lactation^2,3^, while increased feeding also occurs during lactation in humans^1^. In addition to increased hunger, lactating humans sometimes report increased food cravings for palatable high fat and high sugar foods and may exhibit increased attention to cues associated with palatable food^4^. However, the neural circuitry and molecular mechanisms mediating increased palatable food intake and food seeking during lactation are unknown.

Importantly, the lactational period in rodents is a critical period for brain maturation in developing rodent pups, corresponding approximately to a similar developmental timepoint as the third trimester of pregnancy in human fetuses^5–7^. As a result, overconsumption of palatable high fat and high sugar diets during lactation disrupts the normal development of critical hypothalamic and mesolimbic circuitry involved in energy homeostasis and motivation, resulting in an increased risk of the offspring developing metabolic and emotional disturbances later in life in both rodents and humans^6,8–15^. Despite this, the core neural circuitry and molecular mechanisms mediating altered food seeking and consumption of palatable foods during lactation are largely unknown, and a better understanding of these mechanisms is necessary to reduce the risk of metabolic disease in mothers and their children.

Feeding behavior is regulated by homeostatic neural circuitry in the hypothalamus and hindbrain which sense long-term changes in stored energy in the form of fat derived satiety hormones (i.e. leptin), acute satiety signals from the gastro-intestinal tract (i.e. GLP1, PYY, CCK, etc), and stomach derived signals of energy deprivation (i.e. ghrelin)^16–19^. Fasting concomitantly reduces leptin levels and increases ghrelin levels, resulting in the engagement of hypothalamic circuitry which modulate midbrain dopamine circuits to promote food seeking behavior^16,17,19,20^. Metabolic hormones such as ghrelin, leptin, and insulin also act directly on midbrain dopamine neurons to regulate food seeking and palatable food intake in response to energy needs^21–28^. Feeding can also be regulated by mechanisms that do not depend solely on homeostatic need^29^. For example, animals (including humans) will voluntarily consume palatable diets high in fat and sugar in the absence of energy deprivation, an effect that has previously been conceptualized as hedonic feeding^29–31^.

Although many central neurotransmitters and peptides are important for controlling feeding behavior, the neurotransmitter dopamine is essential for controlling multiple aspects of feeding behavior^30^. Rodents lacking dopamine are aphagic and lack the motivation to seek and consume food. These animals ultimately die of starvation unless exogenous dopamine is provided, indicating an essential role for dopamine in food seeking behavior^32^. Moreover, dopamine is particularly critical for promoting motivated food seeking behaviors as dopamine deficient mice will consume food if the food is presented directly in front of the mouse or in its mouth^30,32–34^. These findings suggest that dopamine is particularly important for the initiation of feeding behavior but may not be required for the consummatory phase of feeding. Consistently, extensive literature supports a critical role for dopamine in promoting the incentive salience or wanting of rewards, including food rewards^27,30,31,35,36^. In particular, ventral tegmental area (VTA) dopamine neurons are robustly activated during food seeking in food-deprived animals and by the consumption of palatable diets^30,37^. These neurons ultimately increase food seeking behavior and reinforcement by releasing dopamine in downstream brain structures, including the nucleus accumbens, amygdala, and prefrontal cortex^30,31^. However, the precise role of dopamine in feeding behavior is debated as increases in dopamine levels also dramatically reduce feeding, such as following the administration of dopamine promoting drugs like amphetamines^30,34,38^.

Despite profound hyperphagia during lactation, changes in food motivation and palatable food intake during the lactation period are poorly understood. Furthermore, although mesolimbic dopamine signaling is important for promoting maternal behavior during lactation^39–44^, and alterations in mesolimbic dopamine signaling are associated with post-partum depression risk^44^, the role of dopamine signaling in modulating feeding during lactation is largely unknown. Here, we utilized a combination of pharmacological, chemogenetic, and *in vivo* imaging approaches to characterize the role of VTA dopamine neurons and mesolimbic dopamine signaling in regulating feeding behavior during lactation in mice.

## Results

### Lactating mice exhibit increased food seeking behavior

Although lactation is known to increase feeding in mice, the effect of lactation on food motivation has not been extensively studied. To characterize feeding behavior and food motivation in mice during lactation we utilized home cage feeding devices (feeding experimental device 3, FED3) which allow for the quantification of meal size and meal frequency in a home cage setting^45^ (**Fig. 1A**). These devices can also function as home-cage operant conditioning devices, allowing for the characterization of effortful food seeking in non-lactating and lactating mice. First, we characterized food seeking behavior in *ad libitum* fed non-lactating and lactating mice during a low amount of effort (fixed ratio 1 schedule of reinforcement, FR1). In this assay, 1 correct nose poke (left poke) is required for the mouse to be awarded 1 20mg food pellet, while the incorrect poke (right poke) is not reinforced. As previously reported^2^, during FR1 testing, lactating mice consumed more pellets by increasing their meal size, while meal number was reduced in lactating mice (**Fig. 1B-D**). Lactating mice performed more correct pokes during testing, while incorrect pokes were significantly reduced in lactating animals (**Fig. 1E-F**). As a result, lactating animals were significantly more accurate at poking at the correct nose poke port compared to non-lactating control animals (**Fig. 1G**).

**Figure 1:**
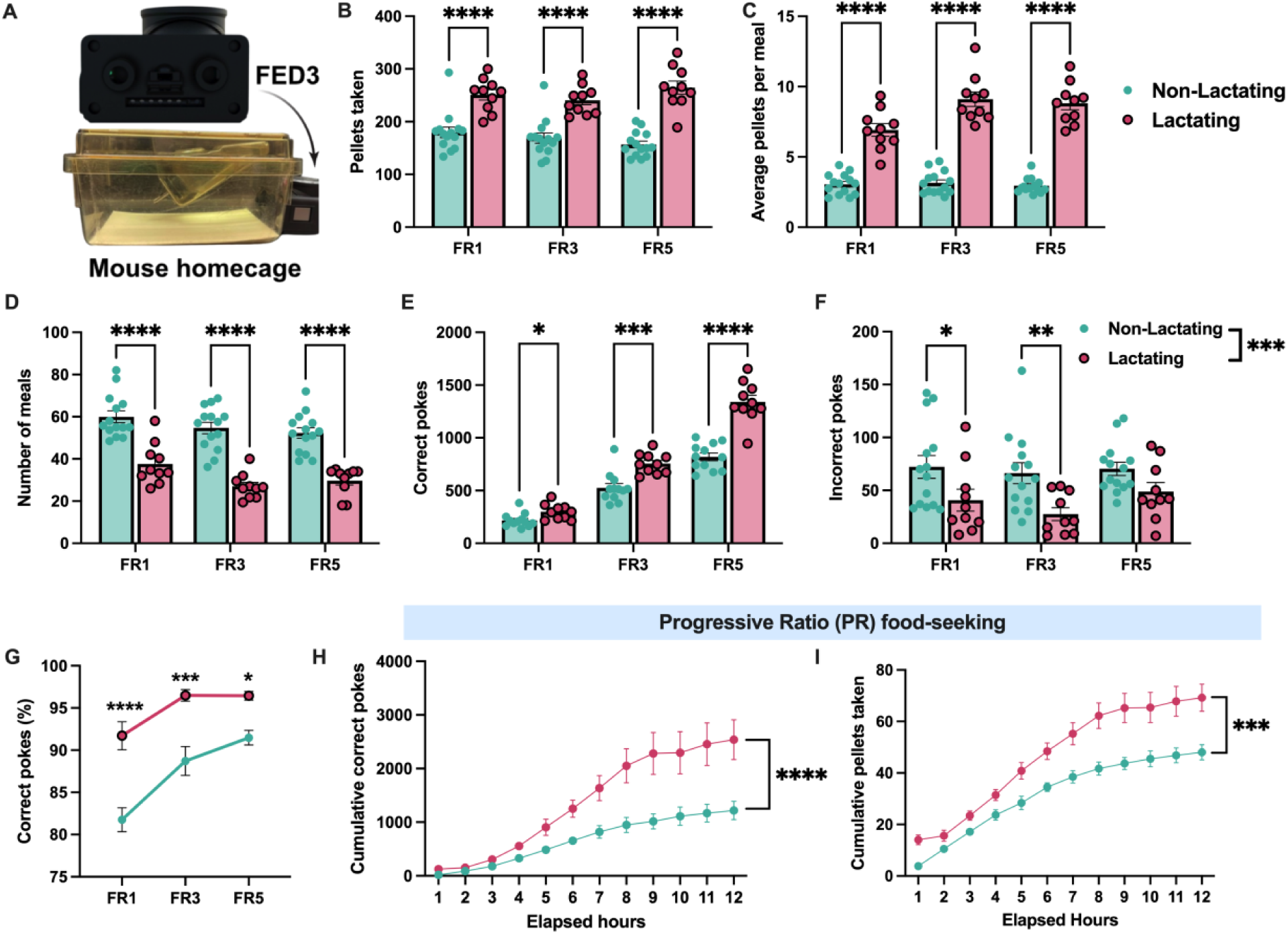
Lactation increases motivated food seeking in mice. (A) Depiction of FED3 device attached to home cage for home-cage operant food seeking behavior. (B and C) Pellets consumed and average number of pellets per meal for non-lactating and lactating mice during fixed-ratio 1 (FR1), fixed ratio 3 (FR3), and fixed ratio 5 (FR5) food seeking assays. (D) Number of meals consumed in non-lactating and lactating mice during operant food seeking assays. (E and F) Correct pokes (E) and incorrect pokes (F) in non-lactating and lactation mice during testing. (G) Percentage of pokes at the correct nose poke port in non-lactating and lactating mice during operant food seeking tasks. (H and I) Cumulative number of nose pokes (H) and pellets consumed (I) during progressive ratio food seeking task in non-lactating and lactating mice. Data represents individual mice. Panels B-G analyzed by 2-way ANOVA with Sidak’s multiple comparison test. Panels H and I analyzed by repeated measures 2-way ANOVA. ns (not significant), *p<0.05, **p<0.01, ***p<0.005, ****p<0.001.

To further test the motivation to seek food in lactating animals, we increased the amount of effort required to obtain food by requiring either 3 correct nose pokes (FR3) or 5 correct nose pokes (FR5) to receive a food reward. Lactating mice exhibited a similar level of hyperphagia during FR3 and FR5 testing as observed during FR1 schedules of reinforcement (**Fig. 1B-D**). Similarly, meal size remained larger while meal number was lower in lactating mice, indicating that increasing the effort required to obtain food does not significantly alter the meal pattern changes associated with lactation (**Fig. 1B-D**). Both non-lactating and lactating mice increased the number of correct pokes when transitioning to higher levels of effortful food seeking (**Fig. 1E**), while the number of incorrect pokes remained relatively stable as the effort required for a food reward increased (**Fig. 1F**). At increasing levels of effort (i.e. FR3 and FR5), lactating mice continued to poke more at the correct poke and less at the incorrect poke compared to non-lactating control mice (**Fig. 1E-F**). Thus, lactating mice exhibit increased food seeking behavior and increased accuracy in food seeking tasks compared to non-lactating animals (**Fig. 1G**).

Since we previously observed increased food seeking behavior in lactating mice at fixed ratio schedules of reinforcement (**Fig. 1B-G**), in new cohorts of mice, we next tested perseverance in food seeking in non-lactating and lactating mice using a progressive ratio reinforcement assay (PR) in *ad libitum* fed mice. In this assay, each successfully retrieved pellet requires an increased number of nose pokes to receive a subsequent food pellet. During PR testing, lactating mice had an approximately 2-fold increase in nose poking relative to non-lactating animals and consumed more pellets than non-lactating animals (**Fig. 1H and I**).

Therefore, lactation increases the effort that mice are willing to provide to obtain food at all schedules of reinforcement.

### Lactating mice exhibit increased palatable food intake

Rodents will voluntarily overconsume palatable food in the absence of hunger (sometimes referred to as hedonic eating). To test if hedonic feeding is also altered in lactation, we measured the consumption of three distinct palatable foods (high fat pellets, sucrose pellets, and peanut butter chips) in *ad libitum* fed non-lactating and lactating mice (**Fig. 2A-C**). All mice were provided access to each of these diets for 10 minutes daily for two days prior to testing to reduce the potential effect of neophobia and to habituate the mice to each diet. During testing, lactating mice consumed more of all three palatable diets relative to non-lactating animals (**Fig. 2A-C**). Since lactating mice generally consume more food than non-lactating animals, we next tested if the relative level of hyperphagia in response to palatable food presentation differs between lactating and non-lactating mice by comparing the number of calories consumed of standard chow and palatable high fat diet 2 hours after presentation. While both non-lactating and lactating mice increased their calorie consumption following acute HFD feeding (**Fig. 2D**), the relative hyperphagia in lactating mice (compared to non-lactating animals) was significantly larger when mice were provided a high fat diet compared to a regular chow diet (**Fig. 2D**).

**Figure 2:**
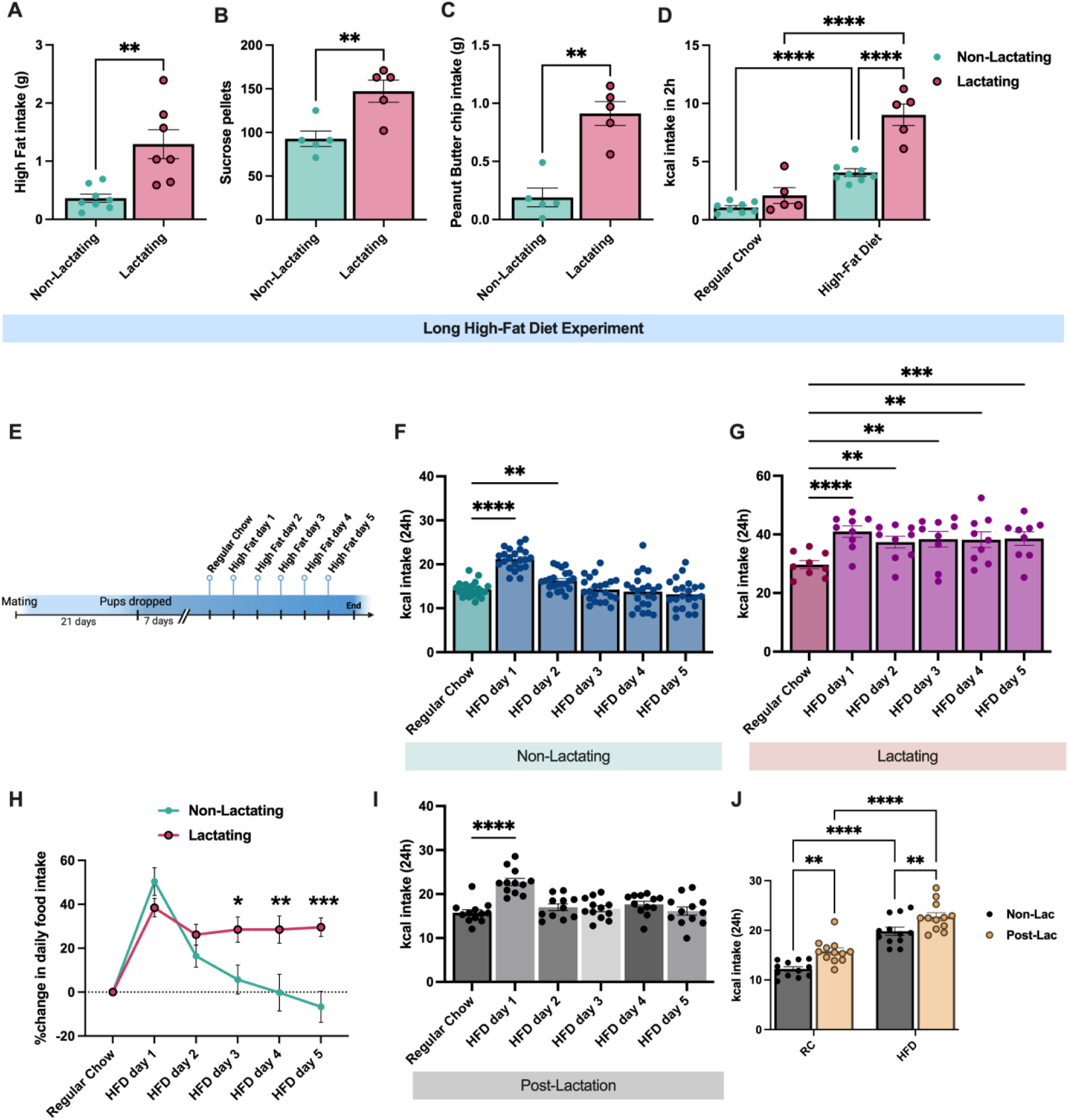
Lactating mice exhibit increased hedonic feeding. (A-C) Intake of high fat diet (A), sucrose pellets (B), and peanut butter chips (C) in *ad libitum* fed non-lactating and lactating mice. (D) Two-hour food intake (in kcal) of non-lactating and lactating mice provided *ad libitum* access to standard chow food or high fat diet. (E) Schematic showing experimental design for daily high fat diet experiments shown in F and G. (F) Daily calorie intake in non-lactating mice following high fat diet administration. (G) Daily calorie intake in lactating mice following high fat diet administration. Individual data points represent individual mice. Data in A-C analyzed by unpaired Student’s t-test. Data in D and J analyzed with repeated measures 2-way ANOVA with Sidak’s multiple comparison test. Data in panels F, G, and I analyzed by repeated measures one-way ANOVA with Dunnett’s post-hoc test. **p<0.01, ***p<0.005, ****p<0.001.

To further characterize the hyperphagic response to high fat diet in lactating mice, in a new cohort of mice, we characterized the feeding response following multiple days of high fat diet feeding (**Fig. 2E**). As expected, non-lactating mice significantly increased their calorie intake following 24 hours of HFD feeding (**Fig. 2F**). This hyperphagia was significantly reduced on the second day of HFD feeding, and caloric intake was undistinguishable from standard chow levels from the 3^rd^ day of HFD feeding until 5 days of HFD feeding (**Fig. 2F**). Like the non-lactating mice, lactating animals significantly increased their caloric intake on the first day of HFD feeding (**Fig. 2G**). However, lactating animals continued to overconsume HFD for 5 consecutive days of HFD feeding, relative to intake of standard chow (**Fig. 2G**). Remarkably, lactating mice did not reduce their hyperphagic response to HFD following multiple days of HFD feeding and continued to consume similar amounts of HFD on the fifth day of HFD access compared to the first day of high fat diet access (**Fig. 2G and H**). The sustained hyperphagia on high fat diet in lactating mice was not due to increasing food intake across the lactational period since lactating mice consumed consistent levels of regular chow across the equivalent time when fed with a regular chow diet (**Extended Data Fig. 1**). Therefore, lactation results in a prolonged hyperphagic response to palatable food compared to non-lactating animals, indicating an increased propensity to overconsume high fat foods during lactation.

Since lactating mice overconsume high fat diet during lactation (**Fig. 2**), in a new cohort of animals, we next tested if this effect persists beyond lactation. Following lactation, we provided high fat diet naïve mice that had previously nursed a litter (10 days post lactation) with daily exposure to regular chow, followed by high fat diet (**Fig. 2I**). In contrast to actively lactating mice (**Fig. 2G**), post-lactation mice exhibited a similar response as virgin mice to daily high fat diet presentation, overconsuming HFD on the first day it is provided and gradually reducing HFD intake in the subsequent days following HFD access (**Fig. 2I**). Interestingly, post-lactating mice consumed more daily calories than virgin age matched mice when provided either regular chow or high fat diet (**Fig. 2J**). Thus, while the relative orexigenic response to HFD is similar between virgin and post-lactating mice, reproductive experience results in elevated calorie intake in the weeks following the lactational period (**Fig. 2J**).

### Dopamine is required for increasing meal size and food seeking in lactating mice

Prior studies indicate a critical role for dopamine signaling in mediating feeding behavior in rodents and humans^30^. However, it is unknown if dopamine is required for the increased food seeking and meal size in lactating rodents. To test for the necessity of dopamine signaling in increasing feeding during lactation, we repeated food seeking assays (FR1) in non-lactating and lactating mice following the administration of the dopamine receptor antagonist flupentixol (0.4mg/kg, i.p.) or saline (**Fig. 3**). As previously described, lactating mice poked more for food and consumed more pellets than non-lactating animals (**Fig. 3A-C**). However, pre-treatment with flupentixol reversed the increased food seeking and food intake in lactating mice to non-lactating levels (**Fig. 3A-C**). Dopamine receptor inhibition also normalized the meal size of lactating mice to non-lactating levels, suggesting that functional dopamine signaling is ultimately required for the increased meal size associated with lactation (**Fig. 3D**). In contrast, acute inhibition of dopamine receptors exerted much more modest effects in non-lactating animals, with no significant differences observed in pellets consumed, nose pokes, or meal size in the hours following flupentixol injections in non-lactating animals (**Fig. 3A, B, D, and F**). Importantly, dopamine receptor antagonism did not alter meal frequency in either groups of mice, indicating that dopamine receptor antagonism specifically reduces feeding by reducing meal size in lactating animals (**Fig. 3E**). Furthermore, it is unlikely that dopamine receptor inhibition significantly disrupted lactation in the dams as pup body weight remained stable following flupentixol injections compared to control saline injections, indicating that pups continued to be nursed following flupentixol injections (**Fig. 3G**).

**Figure 3:**
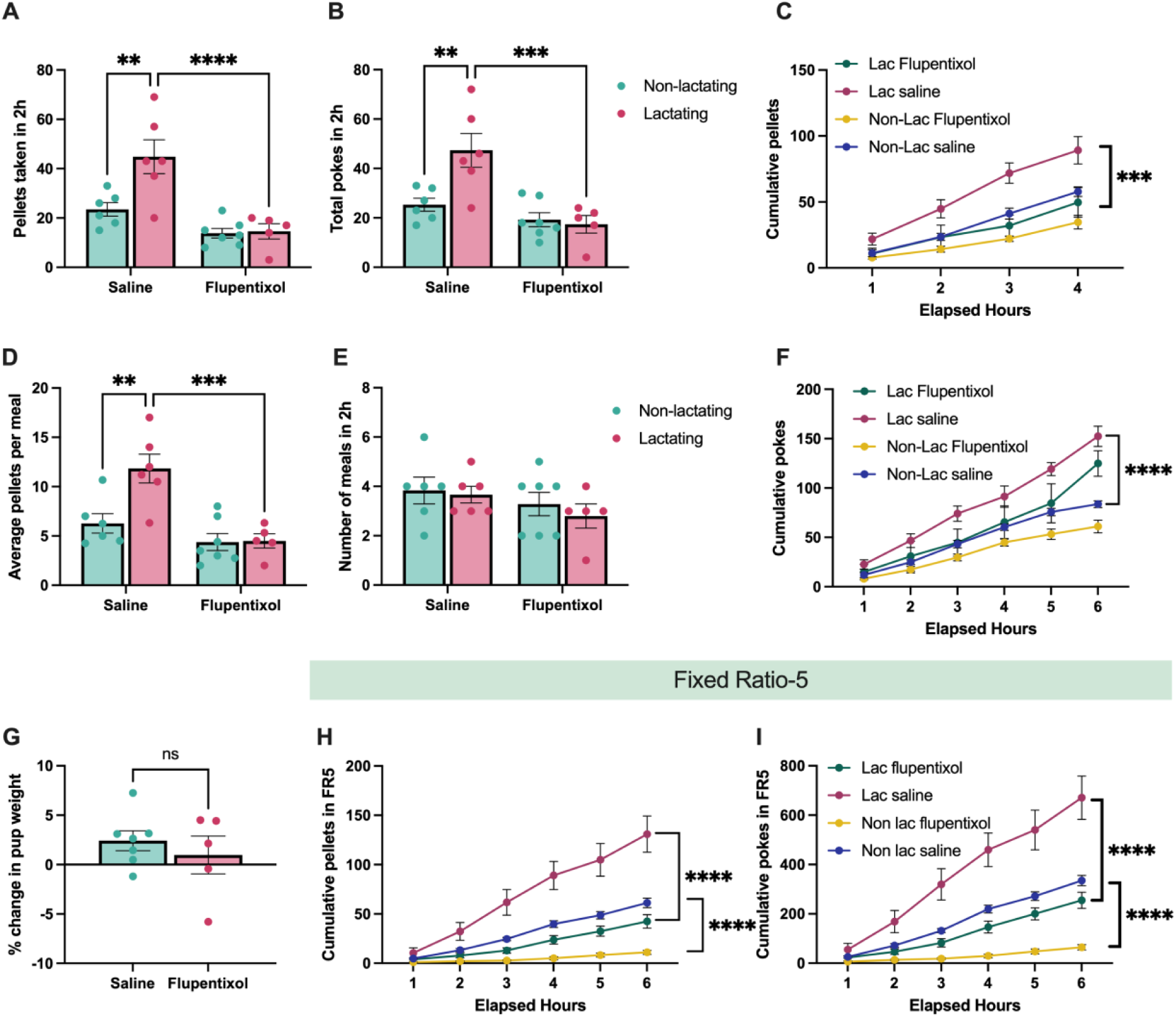
Dopamine transmission is required for the increased food intake, meal size, and food seeking in lactating mice. (A and B) Pellets consumed (A) and total nose pokes (B) in non-lactating and lactating mice injected with saline or flupentixol. (C) Cumulative pellets consumed in non-lactating and lactating mice following i.p. injections of saline or flupentixol. (D and E) Average number of pellets per meal and the average number of meals in two hours following injections of saline or flupentixol in non-lactating or lactating mice. (F) Cumulative number of nose pokes in non-lactating and lactating mice following injections of saline or flupentixol. (G) Change in body weight of the pups following injections of saline or flupentixol. (H and I) Cumulative pellets consumed (H) and cumulative number of nose pokes (I) following injections of saline or flupentixol during fixed ratio 5 (FR5) food seeking tasks. Data points represent individual mice. Panels A, B, D, and E analyzed with 2-way ANOVA with Sidak’s post-hoc test. Panels C, F, H, and I analyzed with repeated measures 2-way ANOVA with Tukey’s post-hoc test. Panel G analyzed with Students unpaired 2-test. ns (not significant), **p<0.01, ***p<0.005, ****p<0.001.

We next tested if dopamine signaling is required for promoting increased food seeking during lactation in conditions of increased effort in non-lactating and lactating mice. In new cohorts of non-lactating and lactating mice we trained mice to nose poke for a food reward at an FR5 schedule of reinforcement (**Fig. 3H and I**). On the testing day, mice were administered either saline or flupentixol and the nose pokes and pellets consumed were measured with FED3 devices. As expected, following saline injections, lactating mice poked significantly more and obtained more pellets than non-lactating mice injected with saline (**Fig. 3H and I**). Pre-treatment with flupentixol reduced both nose pokes and pellets consumed in lactating mice to the levels observed in non-lactating mice following control saline injections (**Fig. 3H and I)**. However, a similar level of inhibition in nose pokes and pellets consumed was observed in both non-lactating and lactating mice (**Fig. 3H and I**). Furthermore, lactating mice continued to poke more for food and obtained more food pellets than non-lactating mice in the presence of dopamine receptor inhibition (**Fig. 3H and I**). Thus, at higher schedules of reinforcement where more work is required to obtain food, global inhibition of dopamine signaling equivalently reduces food seeking in both non-lactating and lactating mice.

### VTA dopamine neurons mediate increased palatable food intake during lactation

Although our prior pharmacological assays indicate an important role for dopamine signaling in regulating feeding behavior during lactation (**Fig. 3**), the specific dopaminergic populations involved are unknown. Given the prevalence of data supporting an important role for ventral tegmental area (VTA) dopamine neurons in regulating food seeking and palatable food intake^30^, we next tested if VTA dopamine activity is required for promoting increased hedonic feeding and food seeking during lactation. To specifically inhibit the activity of VTA dopamine neurons we targeted the chemogenetic inhibitor DREADD hM4Di or control mCherry expressing virus to VTA dopamine neurons in DAT-cre mice (**Fig. 4A and B; Extended Data Fig. 2**). Following viral injections, we mated half of the experimental mice to male animals to induce pregnancy and lactation and tested the feeding behavior of the mice during the lactation period, or the equivalent time in non-lactating animals (**Fig. 4A**). As previously shown, lactating mice consumed more food than non-lactating mice by increasing meal size (**Fig. 4C-E**). To test if VTA dopamine neurons are required for increased regular chow feeding during lactation we injected saline or CNO (1mg/kg, i.p.) to mice expressing either mCherry control virus or hM4Di in VTA dopamine neurons. Chemogenetic inhibition of VTA dopamine neurons did not alter pellet consumption in both non-lactating and lactating mice (**Fig. 4C**). Therefore, VTA dopamine activity is not required for increased regular chow feeding in *ad libitum* fed lactating mice under conditions of limited effort (**Fig. 4C**). As previously reported^46,47^, inhibition of VTA dopamine neurons altered meal structure in non-lactating animals, significantly reducing meal number and non-significantly increasing meal size (**Fig. 4D and E**). Inhibition of VTA dopamine neurons did not significantly alter meal number in lactating mice (although trends towards reduced meal number were observed) and further increased meal size in lactating animals (**Fig. 4D and E**).

**Figure 4:**
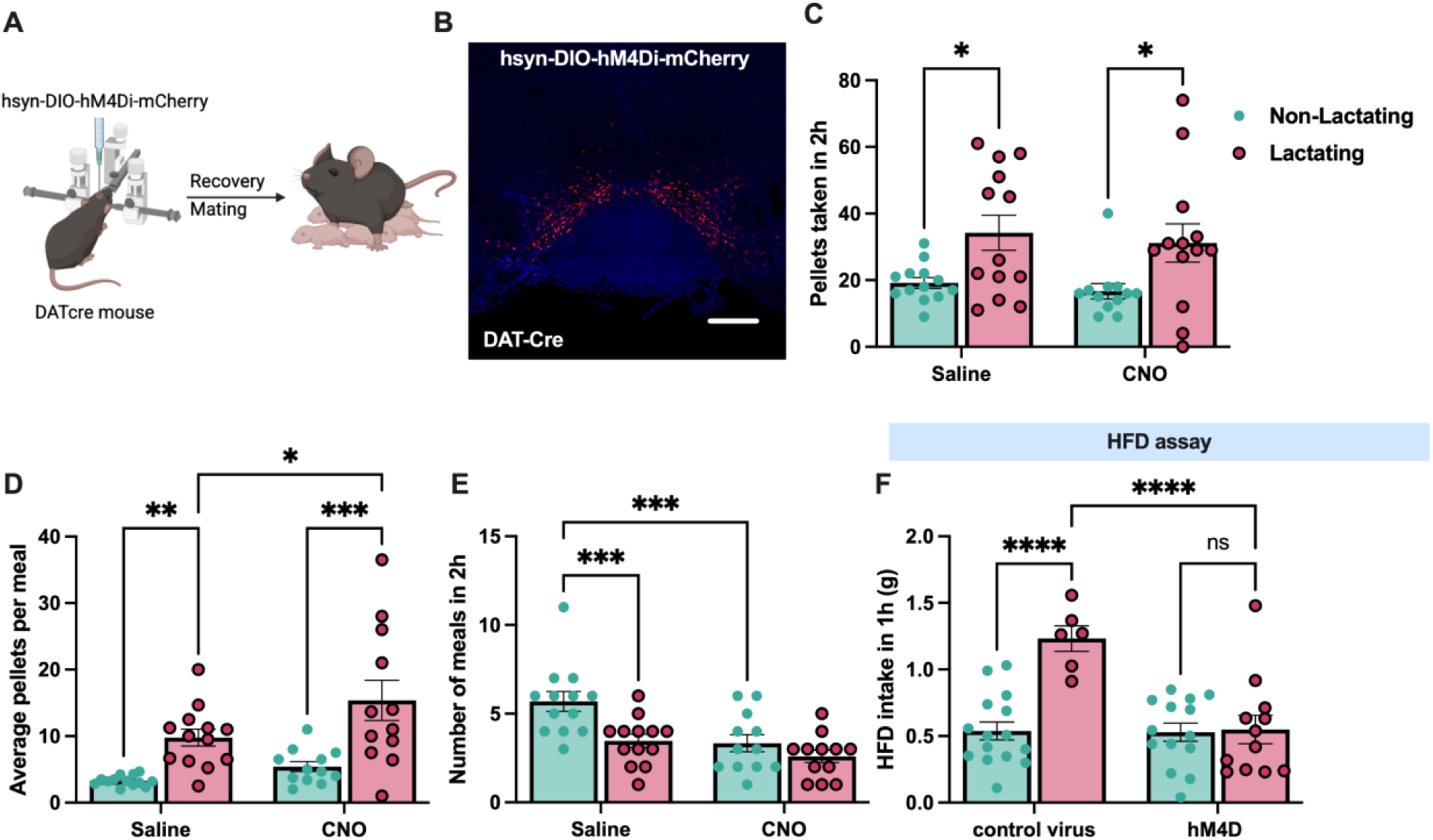
VTA dopamine neurons mediate increased hedonic feeding in lactating mice. (A) Schematic of experimental approach for chemogenetic inhibition of VTA dopamine neurons in non-lactating and lactating mice. (B) Representative image showing hM4Di-mCherry expression in VTA dopamine neurons. (C) Pellets consumed in two hours following injections of saline or CNO (1mg/kg, i.p.) in non-lactating and lactating mice. (D and E) Average number of pellets per meal (D) and average number of meals (E) in non-lactating and lactating mice following injections of saline or CNO. (F) High fat diet consumed in one hour in non-lactating or lactating mice expressing either control mCherry virus or hM4Di expressing virus following injections of CNO (1mg/kg, i.p.). Individual data points represent individual mice. Data in C-F analyzed by Fisher’s LSD test. *p<0.05, **p<0.01, ***p<0.005, ****p<0.001.

No significant effect of CNO administration was detected in pellets consumed, meal number, or meal size in both non-lactating and lactating mice expressing control mCherry virus in VTA dopamine neurons (**Extended Data Fig. 3**). Therefore, although dopamine signaling is required for increased feeding and meal size during lactation (**Fig. 3**), the VTA dopamine neurons are not required for the normal chow hyperphagia and increased meal size observed during lactation in *ad libitum* fed mice. Further work is required to identify the source(s) of dopamine which control meal size during lactation.

VTA dopamine neurons promote increased consumption of rewarding foods and drugs, including high fat diet^30,31^. Since lactating mice exhibited increased palatable food consumption (**Fig. 2**), we hypothesized that VTA dopamine neurons mediate the increased hedonic feeding in lactating mice. To test this hypothesis, we performed acute HFD feeding assays in non-lactating and lactating mice following chemogenetic inhibition of VTA dopamine neurons (**Fig. 4F**).

Consistent with our previous findings (**Fig. 2A-C**), in control mCherry expressing mice, lactating mice consumed significantly more HFD than non-lactating animals following CNO injections (**Fig. 4F**). However, CNO mediated inhibition of VTA dopamine neurons completely reversed the hyperphagia in lactating mice to the levels observed in non-lactating animals (**Fig. 4F**).

Further, inhibition of VTA dopamine neurons did not significantly alter acute HFD intake in non-lactating animals (**Fig. 4F**). Together, these data indicate a specific role for VTA dopamine neurons in mediating the enhanced hedonic feeding associated with lactation, while these neurons are dispensable for normal chow feeding behavior during lactation in *ad libitum* fed mice.

### VTA dopamine neurons mediate increased food seeking during lactation

Although VTA dopamine neurons are not required for promoting food intake in sated animals (**Fig. 4C**), based on prior studies^30,37^, we reasoned that food deprivation may engage VTA dopamine neurons to increase food seeking in a state of negative energy balance. To test if VTA dopamine neurons are required for promoting increased food seeking in food-deprived animals, and for promoting increased food seeking during lactation, we administered CNO to inhibit VTA dopamine neurons following an acute 10 hour fast (**Fig. 5A**). As expected, in control mCherry expressing mice, lactating mice poked more for food and consumed more food than non-lactating animals (**Fig. 5B-C**). In contrast to *ad libitum* fed conditions (**Fig. 4C**), chemogenetic inhibition of VTA dopamine neurons in lactating mice reduced nose pokes and pellets consumed in lactating mice to the levels observed in non-lactating animals (**Fig. 5B-C**). Inhibition of VTA dopamine neurons also significantly reduced nose pokes and food intake in non-lactating mice following an acute fast (**Fig. 5B-C**). As a result, fasted lactating mice continued to poke more for food and consume more food than fasted non-lactating mice during chemogenetic inhibition of VTA dopamine neurons (**Fig. 5B-C**).

**Figure 5:**
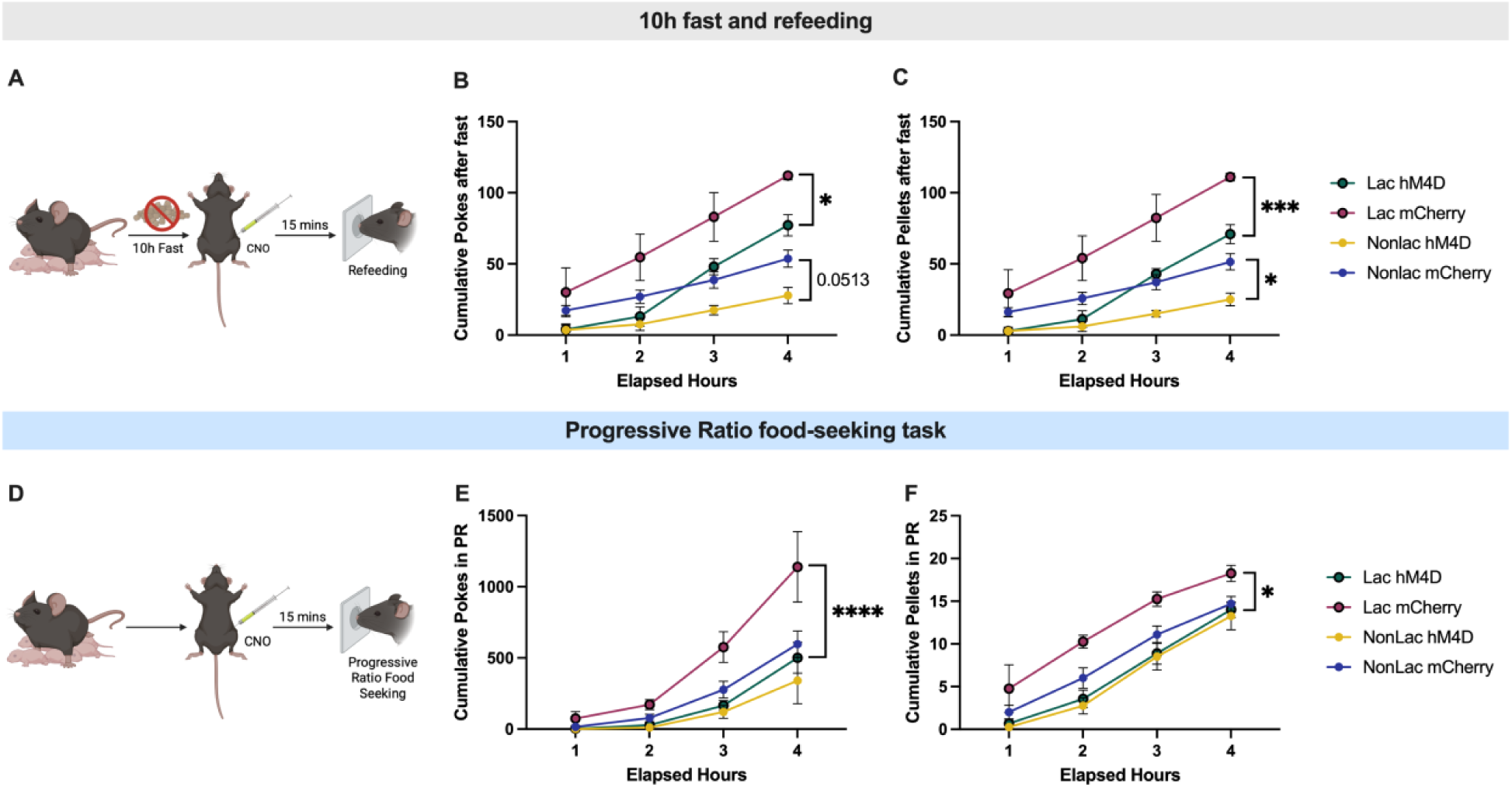
VTA dopamine neurons mediate increased food seeking in lactating mice. (A) Schematic showing experimental design for inhibiting VTA dopamine neurons prior to re-feeding following a ten hour fast. (B and C) Cumulative nose pokes (B) and pellets consumed (C) in non-lactating or lactating mice expressing either control mCherry expressing virus or hM4Di virus in VTA dopamine neurons. All mice were administered CNO (1mg/kg, i.p.) fifteen minutes prior to providing access to food. (D) Schematic showing experimental design for progressive ratio food seeking tasks in *ad libitum* fed non-lactating and lactating mice. (E and F) Cumulative number of nose pokes (E) and cumulative number of pellets consumed (F) following CNO injections in non-lactating or lactating mice expressing either control mCherry virus or hM4Di virus in VTA dopamine neurons. All mice were administered CNO (1mg/kg, i.p.) fifteen minutes prior to PR testing. Data points represent individual mice. Panels analyzed with 2-way ANOVA with Tukey’s post hoc test. *p<0.05, ***p<0.005, ****p<0.001.

VTA dopamine neurons are critically important for driving motivated food seeking in operant feeding assays which require work to obtain food^30,31^, and our prior data show increased food motivation in lactating mice (**Fig. 1**). We therefore next tested if VTA dopamine neurons are required for driving increased food motivation during lactation in *ad libitum* fed mice (**Fig. 5D**). As observed previously, lactating mice expressing control mCherry virus poked more for food and consumed more pellets than non-lactating mice expressing mCherry during progressive ratio food seeking tasks (**Fig. 5E and F**). In contrast, chemogenetic inhibition of VTA dopamine neurons dramatically reduced both nose pokes and pellets consumed in lactating mice (**Fig. 5E and F**). In contrast, inhibition of VTA dopamine neurons did not significantly alter nose pokes or pellets consumed in non-lactating animals (**Fig. 5E and F**).

Thus, following chemogenetic inhibition of VTA dopamine neurons, lactating mice exhibited similar levels of nose pokes and pellets consumed as non-lactating animals, indicating an essential role for VTA dopamine neurons in promoting increased food seeking behavior during lactation.

### Nucleus accumbens dopamine signaling is enhanced during palatable food intake in **lactating mice**

Although our prior pharmacological (**Fig. 3**) and chemogenetic experiments (**Fig. 4 and 5**) confirm an important role for dopamine signaling in promoting feeding behavior during lactation, it is unclear how lactation alters the natural dynamics of dopamine signaling during feeding. Prior work indicates that dopaminergic projections from the ventral tegmental area (VTA) to the nucleus accumbens are engaged during food seeking and the consumption of palatable food, and this increase in dopamine exerts an important role in reinforcing food seeking and promoting palatable food intake^30,31,37,48^. Since we observed increased food motivation and hedonic feeding in lactating mice (**Fig. 1 and 2**), and these effects were attenuated with blockade of dopamine receptors (**Fig. 3**) or inhibition of VTA dopamine neurons (**Fig. 4 and 5**), we hypothesized that lactation enhances the dopaminergic response to food consumption, leading to increased palatable food intake. To test this hypothesis, we performed *in vivo* fiber photometry recordings of dopamine levels during feeding tasks in non-lactating and lactating mice (**Fig. 6A**). Mice were first targeted with the genetically encoded dopamine sensor GRABDA into the nucleus accumbens (NAc)^49^, and a fiber optic cannula was inserted into the NAc for visualizing fluorescent changes with fiber photometry (**Fig. 6A; Extended Data Fig. 4**). In the case of the GRABDA sensor, elevated levels of dopamine are reflected by an increase in green fluorescence, which is detected by the fiber photometry system during awake, freely moving behavior^49^ (**Fig. 6A**).

**Figure 6:**
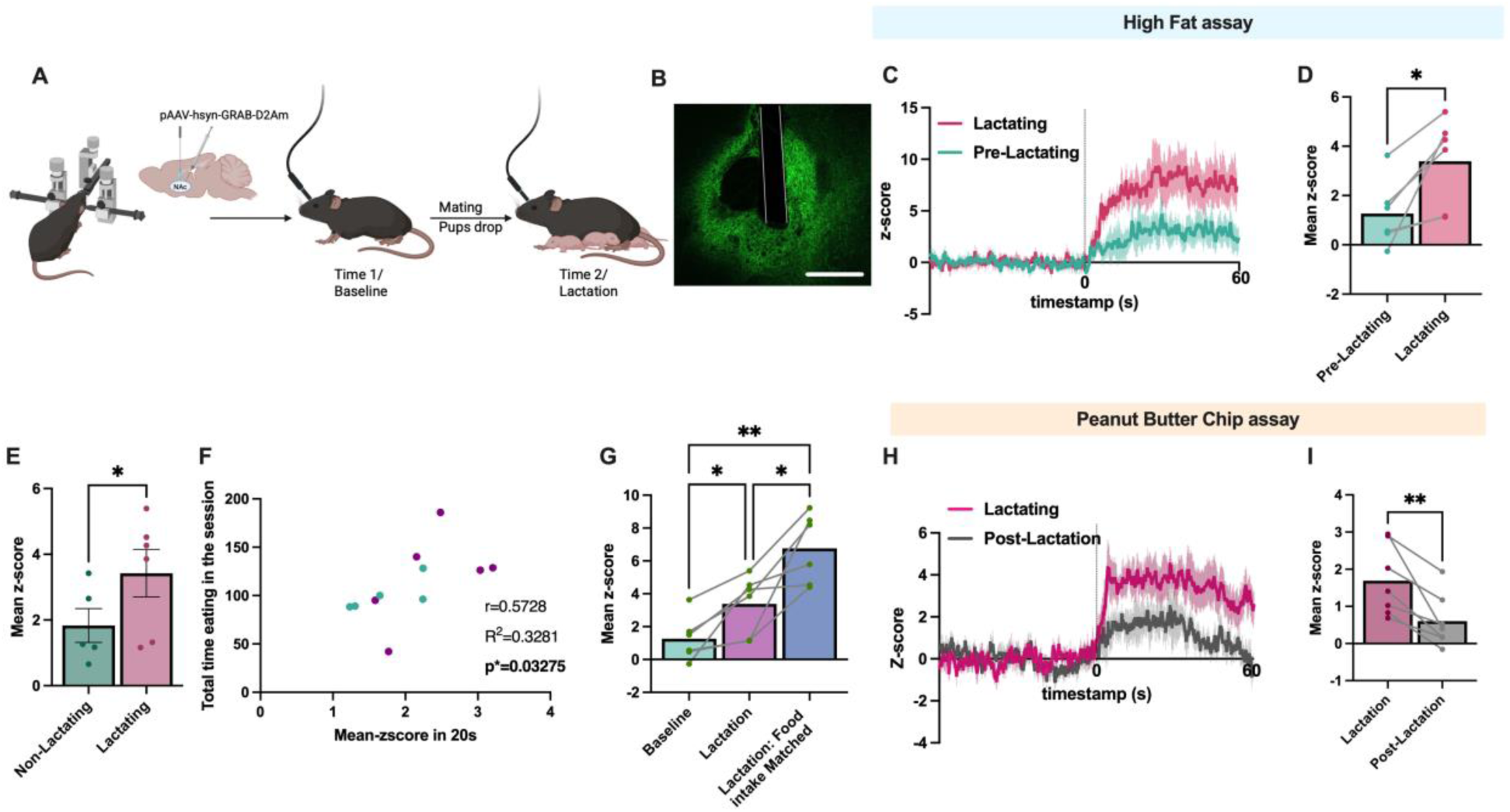
Lactation increases mesolimbic dopamine response during hedonic feeding. (A) Schematic showing experimental approach and timeline for measuring changes in mesolimbic dopamine levels in non-lactating and lactating mice. (B) Representative image showing expression of GRAB dopamine sensor and fiber optic placement in the nucleus accumbens. (C) Average trace of the dopamine response aligned to the consumption of high fat diet during baseline experiments before mating and during the lactational period. (D) Mean Z-score of the average level of dopamine signal following high fat diet consumption in non-lactating and lactating mice for the data shown in C. (E) Mean Z-score showing the change in dopamine levels in the nucleus accumbens following HFD consumption in non-lactating mice and lactating mice on the same testing day. (F) Correlation between the mean Z-score in nucleus accumbens dopamine response following the initial 20 seconds of HFD consumption and the subsequent amount of food consumed during the ten-minute testing session. (G) Change in dopamine levels following HFD consumption in non-lactating, lactating *ad libitum* fed mice, and lactating mice food-intake matched to the levels of non-lactating mice prior to providing high fat diet. (H) Average trace of the dopamine response following peanut butter (PB) chip consumption during the lactational period and one week following lactation. (I) Quantification of the data in H showing the mean change in dopamine signal following PB chip consumption during lactation and following the lactational period.

First, we measured the baseline change in dopamine levels in female mice following the consumption of a palatable high fat diet pellet (**Fig. 6A-C**). As expected, consumption of HFD significantly increased dopamine levels in the nucleus accumbens (**Fig. 6C and D**). Following baseline measurements of the dopamine response to HFD, mice were paired with a male mouse to induce pregnancy and lactation (**Fig. 6A**). A second round of photometry experiments were performed on the same mice during the lactation period, or the equivalent time in mice that did not become pregnant or lactating (**Fig. 6A**). Compared to baseline recordings, dopamine levels were significantly increased in lactating mice during the consumption of HFD (**Fig. 6C and D**). In contrast, HFD consumption increased NAc dopamine levels to a similar amount at each time-point in mice that were not lactating (time-matched control group; **Extended Data Fig. 5**), indicating that these effects are likely not due to changes in fluorescent signal over time.

Consistently, lactating mice exhibited a significantly greater increase in dopamine signal during HFD consumption compared to non-lactating animals on the same test day (i.e. between-subjects comparison, **Fig. 6E**). Since lactating mice consume more high fat diet than non-lactating mice, we also analyzed the increase in NAc dopamine levels in each mouse during the first 20 seconds of HFD consumption, a time point where both non-lactating and lactating mice consumed the same amount of food (**Extended Data Fig. 6**). In the first twenty seconds of consumption, lactating mice had a greater increase in NAc dopamine levels compared to non-lactating mice, indicating that the increase in NAc dopamine signaling in lactating mice is likely not secondary to increased food intake in lactating animals (**Extended Data Fig. 6**).

Furthermore, in both non-lactating and lactating mice, the increase in dopamine signal in the first 20 seconds of consumption correlated with the subsequent amount of food that each mouse consumed in the ten minutes that animals had access to HFD (**Fig. 6F**). To further probe NAc dopamine dynamics in non-lactating and lactating mice, we also recorded NAc dopamine levels following HFD consumption in lactating mice that were matched to the level of food intake observed in non-lactating mice (**Fig. 6G**). This resulted in a more significant increase in dopamine levels in the lactating mice during HFD consumption, compared to *ad libitum* fed lactating mice (**Fig. 6G**). Thus, lactating mice exhibit a greater increase in dopamine levels in the NAc during palatable food consumption compared to non-lactating animals, and this effect is further enhanced when normalizing food intake levels between lactating and non-lactating animals (**Fig. 6G**).

In a new cohort of mice, we next tested if the enhanced dopamine response to palatable food intake during lactation also occurs for other palatable food sources (i.e. peanut butter chips; PB chips). We also tested if this response was reversible following lactation by performing identical experiments on the same mice during lactation and one week after mice stop nursing (**Fig. 6H and I**). As observed for HFD consumption, NAc dopamine levels increased more following PB chip consumption during lactation compared to the post lactation period (**Fig. 6H and I**). In contrast, we observed no significant difference in the NAc dopamine signal in non-lactating mice following PB chip consumption during the equivalent time-periods (**Extended Data Fig. 5**). Therefore, lactation increases nucleus accumbens dopamine levels during palatable food intake, and this effect reverses following the lactational period.

### Mesolimbic dopamine signaling is enhanced during re-feeding in lactating mice

Dopamine levels in the nucleus accumbens (NAc) are also increased during the consumption of food in food-deprived rodents^30,37,48,50^, and this increase in dopamine may regulate the size of ongoing meals and/or the motivation to consume food^30,31^. Given that lactating mice consumed larger meals than non-lactating mice, and this effect was reversed following inhibition of dopamine receptors (**Fig. 3**), we reasoned that lactation may also increase the NAc dopamine response to feeding that is driven by negative energy balance. To test this hypothesis, we food deprived mice for 10 hours and recorded the change in nucleus accumbens dopamine levels following the consumption of food (**Fig. 7A**). As expected, food consumption increased NAc dopamine levels in baseline experiments (**Fig. 7A**; i.e. before the lactational period). As described in prior HFD fiber photometry assays, we repeated this assay during the lactational period, or the equivalent period in mice that did not get pregnant or lactate (**Fig. 7A-B**). In the sixty seconds following food consumption, lactating mice exhibited a 2-3 fold increase in the dopamine response compared to the baseline state (**Fig. 7A-B**). No difference in the dopamine response to re-feeding was detected between the equivalent time-periods in mice that did not become pregnant or lactate (**Extended Data Fig. 5**). Further, lactating mice exhibited a greater increase in dopamine signal during eating compared to non-lactating mice on the same testing day (between-subjects comparison, **Fig. 7C**). Since lactating mice generally consume more food than non-lactating animals following fasting, we further restricted our analysis to the first twenty seconds of food consumption, a time-period where lactating and non-lactating mice consumed the same amount of food (**Extended Data Fig.6**). Consistent with our prior findings, lactating mice exhibited a significantly greater increase in the NAc dopamine response compared to non-lactating animals during the initial 20 seconds of food consumption (**Extended Data Fig. 6**). The increase in nucleus accumbens dopamine response observed in lactating mice during feeding is specific to food consumption, as no significant change in NAc dopamine signal was observed in non-lactating or lactating mice following the presentation of a standard chow pellet to *ad libitum* fed mice (**Fig. 7E**).

**Figure 7:**
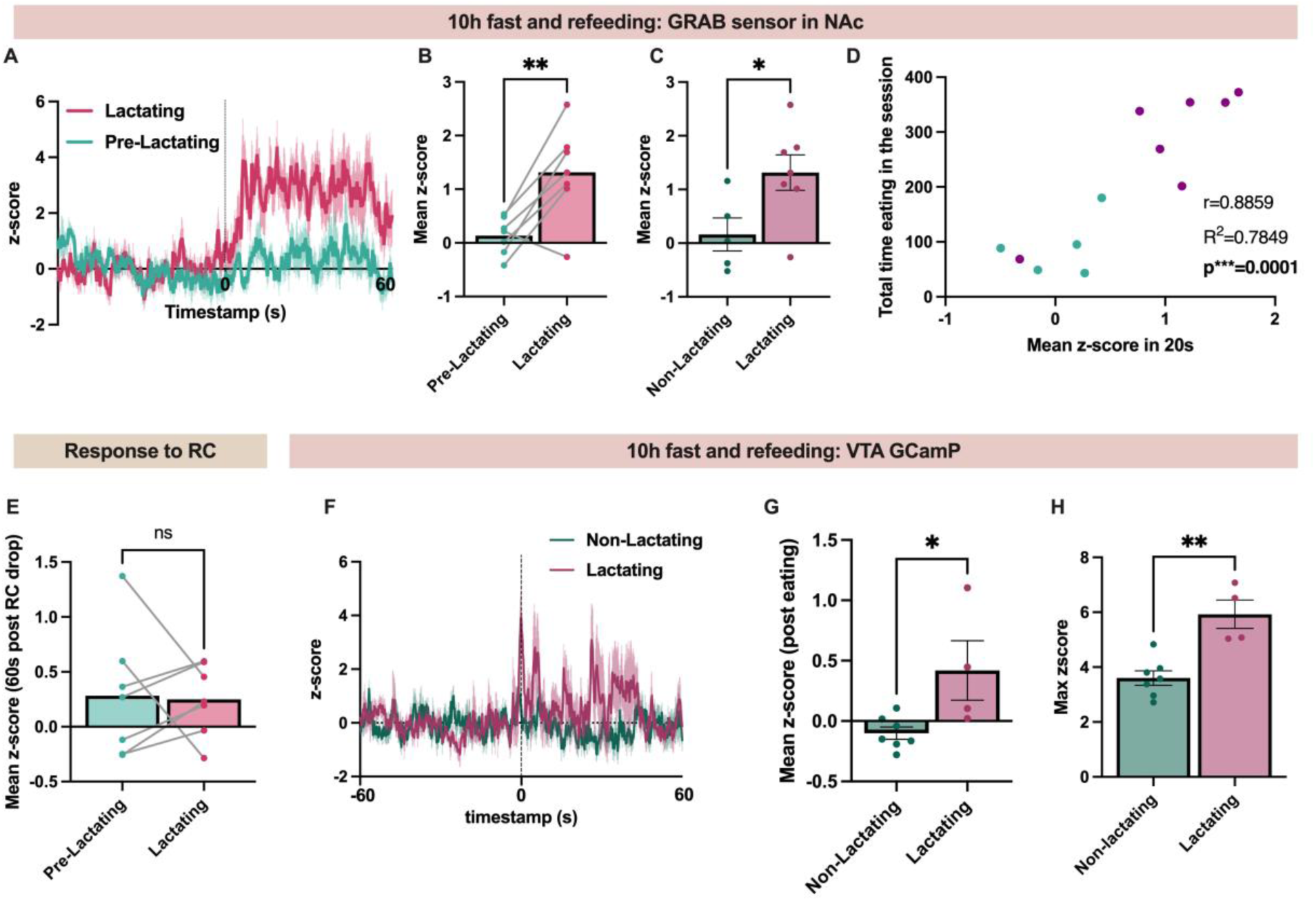
Lactation increases the activity of the mesolimbic dopamine system following re-feeding. (A) Average trace of the dopamine response following the consumption of food after a ten hour fast during baseline conditions prior to mating and during the lactational period. (B) Quantification of the data show in A, comparing the average change in the dopamine signal in the baseline period and during the lactational period. (C) Mean Z-score showing the change in dopamine levels in the nucleus accumbens following food consumption after a ten hour fast in non-lactating and lactating mice during the same testing session. (D) Correlation between the mean z-score of the dopamine response in the first twenty seconds of food consumption and the time eating during the ten-minute testing session. (E) Mean z-score of the dopamine response in the nucleus accumbens in ad libitum fed mice following the presentation of a regular chow food pellet. (F) Average z-score of the calcium activity in VTA dopamine neurons aligned to food consumption in non-lactating and lactating mice. All mice were fasted for ten hours prior to testing. (G and H) Mean (G) and maximum (H) z-score of the VTA dopamine calcium activity following food consumption after a ten hour fast in non-lactating and lactating mice. Graphs A and F show the average trace with the s.e.m. Individual data points in B, C, D, E, G, and H represent individual mice. Data in B and E analyzed with paired Student’s t-test. Data in C, G, and H analyzed with unpaired Student’s t-test. ns (not significant), *p<0.05, **p<0.01.

Next, we tested if the increase in dopamine levels in the nucleus accumbens following feeding is predictive of the amount of food that each mouse will consume. In the first twenty seconds of food consumption, both non-lactating and lactating mice consumed a similar amount of food (**Extended Data Fig. 6**). We thus correlated the change in nucleus accumbens dopamine signal for each mouse in the first twenty seconds of food consumption with the amount of food that each mouse consumed during the ten-minute testing session (**Fig. 7D**).

Interestingly, the initial increase in nucleus accumbens dopamine signal upon eating strongly correlated with the subsequent amount of food consumed during the ten-minute testing session in both non-lactating and lactating mice (**Fig. 7D**). Thus, the initial increase in nucleus accumbens dopamine levels following re-feeding is predictive of the amount of food which will be subsequently consumed.

To confirm if mesolimbic dopamine signaling is enhanced in lactating mice following food deprivation, in a new cohort of mice, we targeted the genetically encoded calcium indicator GCAMP6s to VTA dopamine neurons in DAT-Cre transgenic mice and inserted a fiber optic cannula into the VTA for calcium imaging experiments (**Fig. 7F-H**). Following surgery, mice were randomly split into lactating and non-lactating groups, and the calcium activity of VTA dopamine neurons was recorded following food consumption after a ten hour fast in non-lactating and lactating mice. Like prior results observed while imaging downstream nucleus accumbens dopamine levels in this assay (**Fig. 7A-C**), lactating mice had a greater average change in calcium signal compared to non-lactating mice upon eating (**Fig. 7G**). Furthermore, the maximum change in VTA dopamine activity during initial food consumption was enhanced in lactating mice compared to non-lactating control animals (**Fig. 7H**). Together with prior dopamine measurements in the nucleus accumbens (**Fig. 7A-D**), these results indicate that the mesolimbic dopamine system exhibits increased sensitivity to acute food deprivation in lactating animals.

### Dopaminergic transmission in the NAc regulates meal structure during lactation

VTA dopamine neurons project to multiple downstream brain regions to regulate feeding and motivation^51^. Since we previously observed increased nucleus accumbens dopamine levels following feeding in lactating mice (**Fig. 6 and 7**), we next tested if dopamine signaling in the nucleus accumbens exerts a causal role in regulating feeding behavior in lactating mice. After targeting bilateral guide cannulas to the nucleus accumbens in lactating and non-lactating mice, we repeated FR1 regular chow and HFD feeding assays in *ad libitum* fed mice following intra-nucleus accumbens injections of the dopamine receptor antagonist flupentixol (bilateral, 300nl, 3ug) or aCSF (300nl) (**Fig. 8A-B**). In FR1 feeding assays for standard chow diet, intra-NAc injections of flupentixol significantly reduced total pellet consumption in lactating mice, compared to control aCSF injections (**Fig. 8C-D**). However, the same dose of flupentixol was ineffective at reducing pellet intake in non-lactating animals (**Fig. 8C-D**). As a result, while lactating mice consume significantly more food than non-lactating animals following control aCSF injections, this effect is attenuated following intra-NAc injections of flupentixol (**Fig. 8C and D**). Pharmacological inhibition of DA receptors in NAc did not significantly alter meal size in non-lactating animals, but reduced meal size in lactating mice (**Fig. 8E**). In contrast, like chemogenetic inhibition of VTA dopamine neurons (**Fig. 4**), pharmacological inhibition of NAc dopamine receptors generally reduced the number of meals in both lactating and non-lactating mice (**Fig. 8F**). Thus, NAc dopamine signaling contributes to the increased feeding and meal size in lactating mice, although additional non-NAc dopamine circuits are likely also involved in increasing meal size during lactation.

**Figure 8:**
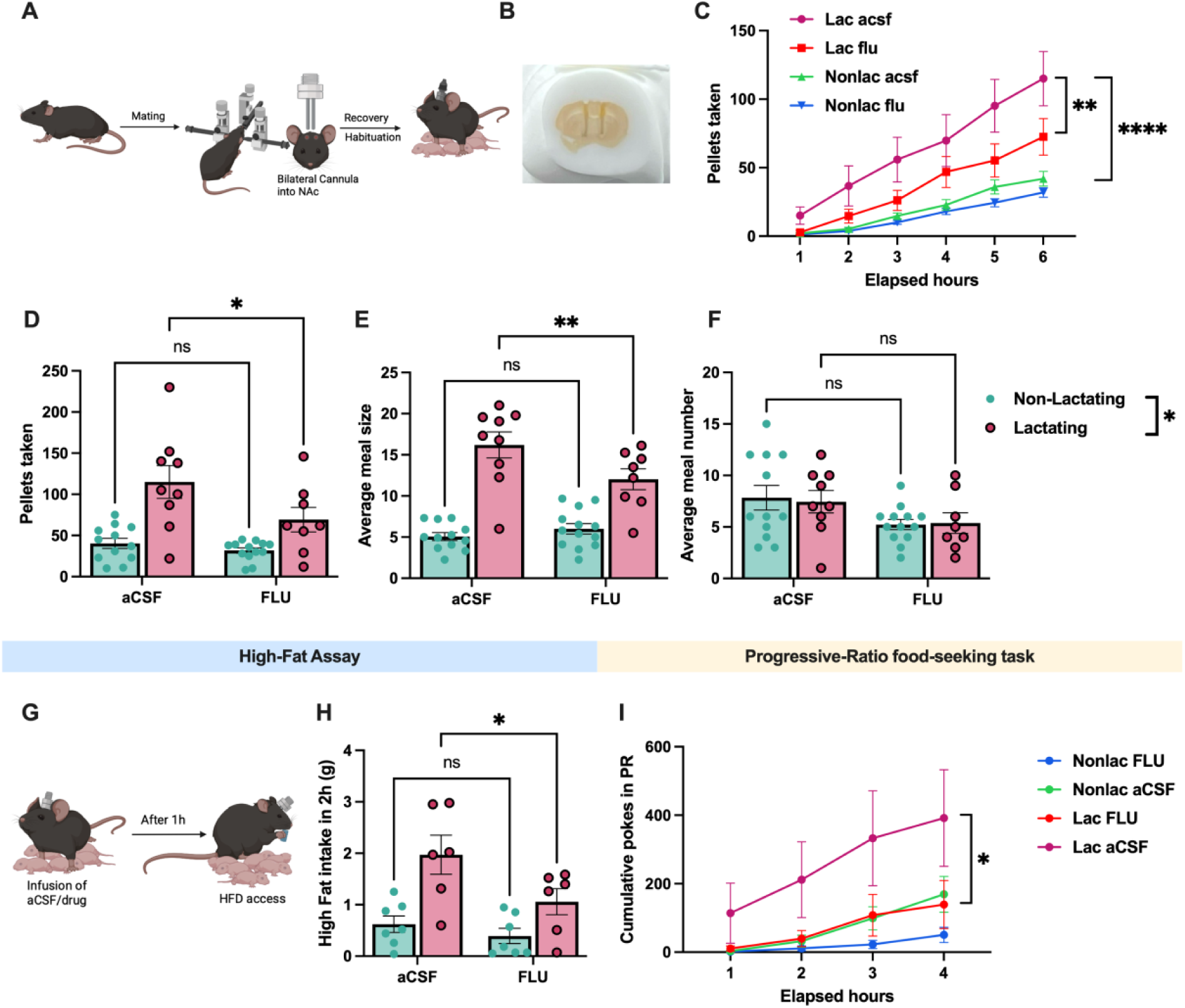
Nucleus accumbens dopamine signaling regulates feeding behavior during lactation in mice. (A and B) Experimental approach (A) and representative image showing cannula track for intra-nuclues accumbens injections (B). (C and D) Pellets consumed following intra-NAc injections of aCSF (300nl) or flupentixol (300nl, 3ug) in lactating and non-lactating animals. (E and F) Average meal size (E) and average number of meals (F) following injections of aCSF or flupentixol in the nucleus accumbent in non-lactating or lactating mice. (G) Schematic showing experimental approach for testing effect of NAc dopamine receptor inhibition on high fat diet intake in non-lactating and lactating mice. (H) High fat diet intake in non-lactating or lactating mice following injections of aCSF or flupentixol. (I) Cumulative nose pokes in non-lactating or lactating mice following injections of aCSF or flupentixol into the nucleus accumbens. Individual data points represent individual mice. Data in C and I analyzed with repeated measures 2-way ANOVA Tukey’s post hoc test. Data in D, E, F, and H analyzed by 2-way ANOVA with Tukey’s post hoc test. *p<0.05, **p<0.01, ****p<0.001.

### Dopaminergic transmission in the NAc regulates palatable food intake during lactation

Inhibition of VTA dopamine neurons reversed the increased hedonic feeding observed in lactating mice (**Fig. 4F**). Since VTA dopamine neurons project densely to nucleus accumbens and this pathway promotes palatable food intake, we next tested if nucleus accumbens dopamine signaling is required for promoting increased hedonic feeding in lactating mice. As expected, lactating mice consumed more high fat diet than non-lactating mice following intra-NAc injections of control aCSF (**Fig. 8G-H**). Inhibition of nucleus accumbens dopamine receptors significantly reduced HFD intake in lactating mice, an effect that was not statistically significant in control non-lactating mice (**Fig. 8H**). Further, HFD consumption in non-lactating and lactating mice were not significantly different in the presence of dopamine receptor inhibition in the nucleus accumbens (**Fig. 8H**). Therefore, functional dopamine receptor signaling in the nucleus accumbens contributes to increased palatable food intake in lactating mice.

### Dopaminergic transmission in the NAc mediates increased food seeking during lactation

Lactating mice poked significantly more for food in progressive ratio food seeking tasks (**Fig. 1**), and this effect required VTA dopamine neurons (**Fig. 5**). Therefore, we tested if downstream dopamine signaling in the nucleus accumbens is required for promoting increased food seeking during lactation by selectively inhibiting NAc dopamine signaling prior to progressive ratio food seeking tasks in non-lactating and lactating mice (**Fig. 8I**). As expected, lactating mice poked significantly more for food than non-lactating mice following intra-NAc injections of aCSF (**Fig. 8I**). Inhibition of nucleus accumbens dopamine receptors drastically reduced nose pokes in lactating mice to levels which were indistinguishable from non-lactating control conditions or non-lactating mice injected with intra-NAc injections of flupentixol (**Fig. 8I**). Although inhibition of dopamine receptors also reduced nose pokes in non-lactating animals, this inhibition was noticeably more modest than observed in lactating mice and failed to reach statistical significance (**Fig. 8I**). Therefore, elevated mesolimbic dopamine signaling from the ventral tegmental area to downstream dopamine receptors in the nucleus accumbens is required for promoting increased food seeking in lactating mice.

## Discussion

Lactating rodents exhibit remarkable hyperphagia, consuming 3-4x more food than control non-lactating animals^1^. Although hyperphagia is less dramatic in humans during lactation, humans also significantly increase feeding during lactation^1^. In addition to increased hunger, lactating women sometimes report increased cravings for palatable food and an increased propensity to consume foods high in fat and sugar^4^. However, the underlying cellular and molecular mechanisms mediating increased food seeking and hedonic feeding during lactation are largely unknown.

Here, we demonstrate that lactating mice drastically increase food seeking behaviors and are more motivated to seek food across multiple food seeking assays (**Fig. 1**). Further, lactating mice exhibit increased consumption of palatable high fat, high sugar foods compared to non-lactating animals (**Fig. 2**). Remarkably, this effect is long-lasting, with lactating mice continuing to overconsume palatable high fat diets for multiple days following access to HFD. In contrast, non-lactating mice appropriately adapt to the increased caloric density of HFD by reducing their voluntary intake of HFD following an initial hyperphagic response (**Fig. 2F and G**). This hyperresponsivity to palatable food intake is specific to the lactational period, as it is not observed in pregnant mice^52^, or following the lactational period (**Fig. 2I**). Given that lactating humans are at an increased risk for developing obesity and metabolic disease^1^, these results suggest that increased food seeking, and elevated hedonic feeding may ultimately contribute to the increased risk for developing metabolic disease in lactating women.

The neurotransmitter dopamine exerts a critical role in promoting food seeking and driving consumption of palatable rewarding foods^30,31,48^. Here, we demonstrate an essential role for dopamine signaling in driving increased meal size, food seeking, and palatable food intake in lactating rodents (**Fig. 3**). Further, although multiple dopamine pathways in the brain regulate feeding behavior^30,32,47,48,53,54^, we provide evidence supporting a critical role for ventral tegmental area (VTA) dopamine neurons in driving both the increased food seeking and the elevated palatable food intake associated with the lactational state (**Fig. 4 and 5**). Consistently, dopaminergic transmission in the nucleus accumbens (a major downstream VTA dopamine projection site) is enhanced in lactating mice following an acute fast and following the consumption of palatable food in the *ad libitum* fed state (**Fig. 6 and 7**). Increased dopamine transmission in the nucleus accumbens likely exerts a causal role in promoting palatable food intake and food seeking behavior during lactation as pharmacological inhibition of dopamine transmission specifically in the NAc significantly reduces palatable food intake and operant food seeking behavior in lactating mice, normalizing the behavior of lactating mice to the levels observed in non-lactating animals (**Fig. 8**). Thus, VTA dopamine projections to the NAc are critical for increasing palatable food intake and food seeking in lactating mice. Importantly, the data presented here does not preclude an additional role for other VTA dopamine projections, such as projections to the basolateral amygdala and prefrontal cortex, in altered feeding behavior during lactation. Further work is required to characterize the role of additional VTA dopamine projections in modulating feeding during lactation, such as the potential role of VTA dopamine projections to the basolateral amygdala in modulating post-ingestive reward during lactation^48^.

In contrast to the important role of VTA dopamine neurons in driving elevated palatable food intake and food seeking behavior in lactating mice (**Fig. 4F**), VTA dopamine neurons are not required for promoting increased consumption of standard chow or promoting increased meal size in lactating mice in conditions requiring minimal work to obtain food (i.e. *ad libitum* fed conditions; **Fig. 4C-E**). For example, inhibition of VTA dopamine neurons did not alter food intake in *ad libitum* fed conditions in either lactating or non-lactating mice (**Fig. 4C**). Recent work suggests that the increased regular chow food intake during lactation is largely mediated by elevated activity of hypothalamic agouti-related protein (AgRP) neurons^2,3^. Together with data presented here, these findings suggest that disparate neural circuitry drive elevated intake of regular chow vs palatable chow during lactation in mice^2,3^. Alternatively, VTA dopamine neurons may selectively regulate feeding during lactation in conditions with elevated reinforcement value or incentive salience associated with feeding^35,36^. Such a hypothesis is consistent with the well documented role for mesolimbic dopamine circuitry in modulating incentive salience or “wanting” behaviors^35,36^. For example, consumption of food in the energy deprived state results in negative reinforcement and thus the incentive salience of food is elevated in this state^55^. Consumption of palatable food in the *ad libitum* fed state results in positive reinforcement and increases the incentive salience of food, a property that requires mesolimbic dopamine circuitry^29,35,56^. In contrast, consumption of regular chow food in the *ad libitum* fed state produces limited reinforcement value and “wanting” for food is low in this state. Thus, VTA dopamine neurons may be less critical for promoting feeding in this context. Further work is required to dissect the specific function of mesolimbic dopamine circuitry in regulating reinforcement associated with feeding during lactation.

Although inhibition of VTA dopamine neurons did not alter regular chow food intake in lactating and non-lactating mice, inhibition of these neurons generally increased meal size and reduced meal number in both non-lactating and lactating mice (**Fig. 4D-E**). This data is consistent with recent reports suggesting a role for VTA dopamine neurons in regulating the size of ongoing meals and/or the propensity to initiate a meal^46,47^. Additional non-VTA dopamine pathways likely contribute to increased feeding and meal size during lactation in mice since global inhibition of dopamine signaling reduces standard chow intake and meal size in lactating mice (**Fig. 3**). It is important to note that many additional non-VTA dopaminergic pathways are known to regulate feeding behavior, such as the dopaminergic neurons in the arcuate nucleus^53^, zona incerta^47^, nucleus of the solitary tract^54^, and substantia nigra^32,33^, and these cell populations were presumably targeted by peripheral inhibition of dopamine receptors. Further work is required to fully characterize the role of each of these dopaminergic pathways in promoting food intake and meal size in lactating animals.

Notably, inhibition of nucleus accumbens dopamine receptors reduced regular chow food intake in lactating mice (**Fig. 8**), while chemogenetic inhibition of VTA dopamine neurons did not produce this effect (**Fig. 4**). This suggests that other dopaminergic inputs to the nucleus accumbens (other than the VTA) and/or presynaptic dopamine receptors localized to the nucleus accumbens may also be involved in regulating standard chow food consumption and meal size during lactation. Alternatively, this apparent discrepancy may result from differences in the two technical approaches, such as pharmacokinetic differences between chemogenetics and pharmacology or a more pronounced physiological effect on nucleus accumbens neurons following local blockade of dopamine receptors (compared to chemogenetic inhibition of VTA DA neurons). Thus, although accumbal dopamine release contributes to feeding behavior changes during lactation, further work is required to precisely characterize the role of various dopaminergic inputs to NAc in controlling food intake and meal size during lactation.

Additionally, future studies should evaluate the downstream dopamine receptors in NAc (i.e. D1 vs D2 dopamine receptors), and the specific cell types in the nucleus accumbens which mediate altered feeding during lactation.

Although the current data indicates a critical role for mesolimbic dopamine circuitry in promoting increased palatable food intake and food seeking during lactation, the neurophysiological and hormonal mechanism(s) mediating these effects are unknown.

Importantly, the lactational period is associated with dramatic changes in neuroendocrine and reproductive hormones, including increased release of oxytocin and prolactin, and suppression of estradiol levels^1,57^ Click or tap here to enter text.. Prior work indicates that these hormones modulate the activity of the mesolimbic dopamine circuit to promote maternal behavior during lactation^40–44^, and similar mechanisms may also rewire the mesolimbic circuitry to facilitate the changes in feeding outlined here. Future work is required to map the neuroendocrine circuitry linking lactation to increased feeding behavior, and to establish how lactation alters the neurophysiological properties of VTA dopamine neurons.

In addition to the neuroendocrine changes associated with lactation, the energetic demands of milk production and maternal care result in a state of negative energy balance during lactation^1,57^. This increase in energetic demands is coupled to elevated activity of hypothalamic agouti-related peptide (AgRP) neurons, which promote standard chow feeding during lactation^2,3^. Interestingly, elevated AgRP neuron activity modulates mesolimbic dopamine circuitry to promote palatable food intake in conditions of negative energy balance, such as following caloric restriction^48,50,58^. For example, chemogenetic activation of AgRP neurons enhances the mesolimbic dopamine response to palatable food^50^, and this effect is hypothesized to drive increased food intake. Given that the lactational state is a condition of negative energy balance and increases AgRP neuron activity^2,3^, functional interactions between the hypothalamic melanocortin system and dopamine signaling are likely important for promoting increased food seeking and palatable food intake during lactation in mice. Future work is warranted to characterize the mechanisms mediating functional interactions between hypothalamic AgRP neurons and dopaminergic transmission in regulating feeding during lactation.

In conclusion, we outline an essential role for mesolimbic dopamine circuitry in driving increased food seeking and palatable food intake in lactating mice, providing a cellular mechanism mediating food cravings for palatable food and increased food seeking behaviors during the lactational state.

## Methods

### Animals

Female C57/BL6J mice (Jax#000664) or DAT-Cre mice (Jax# 020080) were used for all experiments. DAT-Cre female mice for experiments were obtained by breeding heterozygous Cre mice to C57/BL6J mice purchased from the Jackson Laboratory. Weanlings were genotyped and Cre+ mice were used for chemogenetic and GCAMP fiber photometry experiments. All mice were group caged in 12h light/dark cycle, temperature (70F) and humidity-controlled rooms. Food and water were provided *ad libitum* except during specific experimental conditions, as noted in the text. All experimental procedures were approved by University of Illinois Institutional Animal Care and Use Committee (IACUC).

To generate experimental mice for all experiments (i.e. lactating animals), 8–14-week-old female mice were mated with reproductively experienced male mouse for 5 days. After the male mouse was removed from the cage, pregnancy progression was monitored until the lactational period. Control groups included mated females that did not become pregnant as well as virgin females that were group-housed with a second female mouse for 5 days. Experiments were conducted between lactation day 5 and lactation day 18.

### Feeding behavioral assays

For most experiments, food intake was quantified using Feeding Experimentation Devices v3 (FED3; Open Ephys), which were fit into modified home cages (Figure 1A). Each FED3 unit consisted of two nose-poke ports, positioned on the left and right side of a central pellet dispenser. The left port was designated as the correct nose poke, whereas the right port was designated as incorrect. Nose pokes on the incorrect port did not result in pellet delivery. Operant schedules of reinforcement required a defined number of correct nose pokes for the delivery of a single pellet. Under the fixed ratio 1 (FR1) schedule, one correct nose poke resulted in pellet delivery; under fixed ratio 3 (FR3), three consecutive correct nose pokes were required; and under fixed ratio 5 (FR5), five consecutive correct nose pokes were required. For the progressive ratio (PR) schedule, the number of nose pokes required increased progressively with each successive pellet. Progressive ratio schedule utilized a linear progression model for figure 1 (i.e. 1 poke, 2 pokes, 3 pokes, 4 pokes, etc). Progressive ratio schedule was as follows for chemogenetic and flupentixol injection experiments: 1, 2, 4, 6, 9, 15, 32, as outlined in a previously publication^30,32–34,58^. Breakpoint was calculated as the number of pokes by the mouse before it stops poking for at least 2 hours.

Meal size was defined as the number of pellets consumed consecutively, separated by an inter-pellet interval of ≤5 minutes. Meal number was defined as the total number of such meals occurring within a specified time interval. Accuracy was calculated as the percentage of total nose pokes that were correct divided by the total number of pokes.

### Hedonic Feeding Assays

For acute hedonic feeding assays, *ad libitum*–fed mice were given access to either a high-fat diet (60% HFD, Research Diets) or peanut butter chips (Reese’s) for a 1-hour period. To habituate mice to the palatable food, animals were given access to the diet (high fat or peanut butter chips) for 10 minutes daily for two days prior to the experimental session. Sucrose intake was measured using the FED3 devices. FEDs were loaded with sucrose pellets (Bio-Serv) and set to a fixed ratio 1 (FR1) schedule. Mice had continuous access to standard chow in the home cage, and sucrose intake was quantified over a 24-hour period.

### Long High Fat Diet experiment

After recording 24-hour body weight and regular chow intake for 2 days, mice were given access to only high fat diet pellets in the cage food hopper for 5 days. During this time, daily body weight and food intake measurements were taken at start of the dark cycle by manually measuring the amount of food left in each cage and comparing this to the amount of food provided to each mouse on the previous day. Cages were changed daily during testing, and mice were provided with new HFD each day.

### IP Flupentixol injections

Cis-Z-Flupentixol (MedChemExpress; HY-15856) was suspended in 0.9% saline to make stock solution. For all experiments, mice were administered 0.04mg/kg dose intraperitoneally at the start of dark cycle. The mice were first habituated to saline injections and the FEDs prior to start of the experiments for at least two days before administering flupentixol. For feeding assays, mice were administered either saline or flupentixol in a randomized and counterbalanced fashion such that half of the mice received flupentixol and the other half received saline. Treatment groups were flipped on the following day, with each mouse receiving the opposite treatment condition. To determine if flupentixol injections altered the ability of the dams to nurse their pups, the body weight of the pups was measured immediately prior and 6 hours after injections of saline or flupentixol.

### Stereotaxic surgery

Stereotaxic surgeries were performed as previously described^2^. Mice were anesthetized with isoflurane (1–2% in oxygen) and placed in a stereotaxic frame (Kopf Instruments) with continuous isoflurane and oxygen delivery. Preoperative analgesia was provided with carprofen (5 mg/kg, s.c.), which was continued for two days postoperatively.

After confirming deep anesthesia, the scalp was shaved, and incision was made to expose the skull. A small burrhole was made directly above the viral injection site, and viral vectors were delivered to the ventral tegmental area (VTA) or nucleus accumbens (NAc) using a pulled glass micropipette connected to a microinjector (Ronal tools). Injection coordinates (from bregma) for VTA targeting were: A/P = −2.8 mm and −3.1 mm, M/L = ±0.35 mm, D/V = −4.2 mm (from brain surface). AAV-syn-flex-GCAMP6s or AAV-hM4Di/mCherry (120 nL/site) were injected into the VTA, and GRAB-DA sensors were injected into the NAc at the following coordinates: A/P = 1.2 mm, M/L = 1.0 mm, D/V = 4.1mm, 3.8mm and 3.5mm. Injections of hM4Di-mCherry were bilateral, whereas GCaMP6s and GRAB-DA injections were unilateral.

Viral solutions were infused at a speed of 60nl per minute and the pipette was left in place for 5 minutes after infusion before withdrawing to minimize backflow and leakage of vírus outside the desired target structure.

For fiber photometry experiments, an optical fiber (200 µm core, RWD Biosciences) was implanted 0.2 mm above the viral injection site (VTA or NAc) during the same surgery. The implant was secured to the skull with dental cement (C&B Metabond). Mice were singly housed and allowed to recover for at least three weeks before experiments.

### Viral vectors

Adeno-associated viral vectors (AAV) that were used in this study included Cre-dependent GCAMP6s (AAV5-Syn-Flex-GCAMP6s-WPRE-SV40; #100845), Cre-dependent hM4Di (AAV5-hsyn-DIO-hM4Di-mCherry; #44362), Cre-dependent mCherry control virus (AAV5-hsyn-DIO-mCherry; #50459) or Cre-dependent GFP control virus (AAV5-hsyn-DIO-EGFP; #50457), and GRAB dopamine sensor virus (AAV9-hsyn-GRAB-DA2m; #140553). All viruses were purchased from addgene and were injected into the brain at stock concentrations (> 1x10^12 vg/mL).

### Chemogenetic experiments

For chemogenetic feeding assays, virally injected female DAT-Cre mice were mated to reproductively experienced males for 5 days. Control mice were paired with another female mouse for 5 days. 5 days after birth, dams were habituated to the saline injections and given 10 minutes access to high fat diet for two days. For high fat diet feeding assays, on the day of the experiment, all mice (mCherry control and hM4Di expressing mice) were injected intraperitoneally with Clozapine-N-Oxide (CNO; Enzo Life Sciences) and after a 15-minute wait period, were given high-fat diet for 2 hours, and food intake was measured after 2 hours of access to high fat diet.

Regular chow feeding assays (i.e. FR1 feeding assays) with chemogenetic manipulation of VTA dopamine neurons were performed using a repeated measures analysis approach. FR1 feeding assays were performed once all mice poked at the correct nose port at least 70% of the time (2-3 days of habituating). During habituation, all mice were administered saline daily (i.p. 200ul) to habituate mice to handling and i.p. injections. Following baseline saline injections, half of the mice were injected with CNO (1mg/kg, i.p., 200ul of saline) and the other half received 0.9% saline (i.p., 200ul) 15 mins before the start of the dark cycle. The next day, the groups were flipped, and data was collected on FR1 mode of the FEDs to quantify pellets consumed, meal size, and meal number following saline or CNO injections for each mouse. Following FR1 feeding assays, mice were sequentially advanced to FR3 and FR5 schedules of reinforcement until each mouse poked at least 70% correctly and the correct nose poke. During this period, all mice continued to receive saline injections daily. For the Progressive Ratio experiment, all mice received CNO injection 15 minutes prior to the FEDs being switched to PR mode at the start of dark cycle and data was compared between control mCherry/GFP and hM4Di expressing mice (i.e. between subjects comparison). High fat diet feeding and progression ratio assays were performed using a between-subjects analysis approach due to the potential for order effects associated with these assays, while FR1 regular chow feeding assays were analyzed by using a repeated measures paired analysis comparing the behavior of each mouse when administered saline vs CNO.

In a separate cohort of mice, to test the effect of VTA dopamine neuron inhibition in fasted mice, mice were fasted for 10 hours during the day and refed at the start of dark cycle. 15 minutes before refeeding, all mice were administered CNO and pellets consumed and nose pokes were measured using FED3 devices.

### Fiber photometry

Fiber photometry recordings were conducted using a multi-wavelength fiber photometry system (Plexon multi-wavelength fiber photometry system, 8-61-A-07-A). Mice were connected to the system via a fiber optic patch cord (Plexon, 08-60-A-04-C) coupled to the implanted cannula (RWD biosciences, 200um fiber optic cannula, 0.50 NA) using mating sleeves (Plexon). Excitation light at 465 nm (GRAB-DA/GCaMP6s signal) and 410 nm (isosbestic control) was delivered and emitted fluorescence was collected. Fluorescence signals were processed and analyzed using custom R scripts as previously described^2^.

For GRAB dopamien experiments, the signal trace of all the mice in response to high-fat diet (HFD) pellet was first recorded in an initial session to verify viral expression. Only animals that had changes in 465 nm fluorescence during HFD consumption were included in subsequent experiments. Mice without signal were considered misses and were not included in subsequent experiments. Approximately 50% of virally targeted and fiber implanted mice had measurable signal and were included in experiments.

Following baseline verification of signal, for GRABDA fiber photometry experiments, mice were recorded both prior to mating and during lactation (days 7–14 postpartum, or the corresponding time period in non-pregnant control animals). In the first experiment, dopamine dynamics were recorded during chow consumption under fed and fasted conditions. For the fed condition, recordings consisted of a 5-minute baseline period followed by 5 minutes of chow access. The procedure was repeated after a 10-hour fast (7:00 a.m.–5:00 p.m.) using identical baseline and recording periods.

In the second experiment, dopamine responses to palatable HFD consumption were assessed. Mice were habituated to the HFD by receiving 10 minutes of home-cage access on two consecutive days prior to testing. On the experimental day (between 10:00 a.m. and 4:00 p.m.), mice were connected to the photometry system and recorded for a 5-minute baseline period followed by 10 minutes of HFD access. For both re-feeding following standard chow and HFD access experiments, dopamine or calcium traces were aligned to the start of eating. For the food intake matched lactating mice, the amount of food consummed by non-lactating mice was measured from 10am-4pm (lights on period). Lactating mice were provided with a premeasured amount of food equivalente to the average amount of food consummed by non-lactating animals between 10am-4pm, and fiber photometry experiments were performed on the lactating mice as described above after 6 hours of access to non-lactating levels of food.

Behavioral events were manually logged using a Plexon event input generator. Signal changes were quantified by comparing fluorescence during a 60-second window following eating onset to the 60-second baseline preceding each event. Within-subject comparisons were performed to assess how lactation modulated nucleus accumbens dopamine responses to chow and HFD consumption (i.e. comparing baseline response to response during the lactation period). Between-subjects experiments were also performed to compare the change in dopamine signal between non-lactating and lactation mice following consumption of chow or HFD on the same testing day. For VTA calcium imaging experiments, control non-lactating mice exhibited significant signal decay when comparing the baseline measurement to the lactational period. For this reason, we only analyzed the change in VTA calcium activity in non-lactating vs lactating mice on the same test day during the lactational period. GRABDA dynamics were stable in control mice when comparing the baseline period to the lactational period. Thus, for GRABDA analysis we compared both the baseline period to the lactational period in the same mice (i.e. repeated measures design), and the change in DA signal in non-lactating to lactating mice on the same day during the same testing session (i.e. between subjects experimental design).

### Cannula implantations and microinfusions

Bilateral cannula (RWD) were implanted during stereotaxic surgery into WT lactating mice on day 3 of lactation. The surgery was performed as described above for viral injection surgeries, with cannula’s targeting the nucleus accumbens at the same coordinates utilized for GRABDA fiber photometry experiments. After a week of recovery, the mice were habituated to aCSF (300nl) infusions daily along with the FEDs. The infusion volume was 300nl and infused over 60 seconds using an infusion pump (RWD Biosciences) connected to a Hamilton syringe. After the mice reached 70% accuracy on the FEDs, we infused half the mice with 3ng of flupentixol (in 300nl aCSF) and the other half received aCSF at the start of dark cycle. The next day, the groups were flipped, and data was collected for the mice on FR1. Pellets consumed, meal number, and meal size were compared between the same mice following either aCSF or flupentixol injections (repeated measured design). Following a two-day rest period, where all mice received daily 10-minute access to HFD and acsf infusions, half the mice received 3ng of flupentixol and other other received acsf. An hour after the infusions, mice were given access to the HFD for 2 hours and high fat diet intake was measured manually as described above. The following day, all mice were subsequently trained on the FEDs on FR3 and FR5 along with acsf infusions daily. On the experimental day, the mice were infused with either flupentixol or acsf and the FEDs were set to PR at the start of dark cycle. For acute HFD and progressive ratio assays, to reduce the potential risk of order effects associated with performing these assays multiple times, food intake and food seeking behaviors were compared between different mice on the same testing day (i.e. mice injected with aCSF or flupentixol).

### Statistical Analysis and data analysis

Specific statistical tests are outlined in the figure legends. Data that was normally distributed was analyzed with parametric statistical tests, while data that was not normally distributed was analyzed with non-parametric tests. All mice with viral injections or fiber optic/cannula implants were perfused following experiments and the location of viral expression and fiber placement was confirmed for all mice (see Extended Data Figures). Only mice with accurate viral expression in the target of interest and fiber optic/cannula placments in the target of interest were included in the data analysis. Data was analyzed using Graphpad Prism.

## Acknowledgments

We thank all members of the Sweeney lab for helpful feedback on this study. This study was funded with support from the University of Illinois and the National Institute of Health (NIH R01HD113522 to PS)

---

**Extended Data Figure 1.**
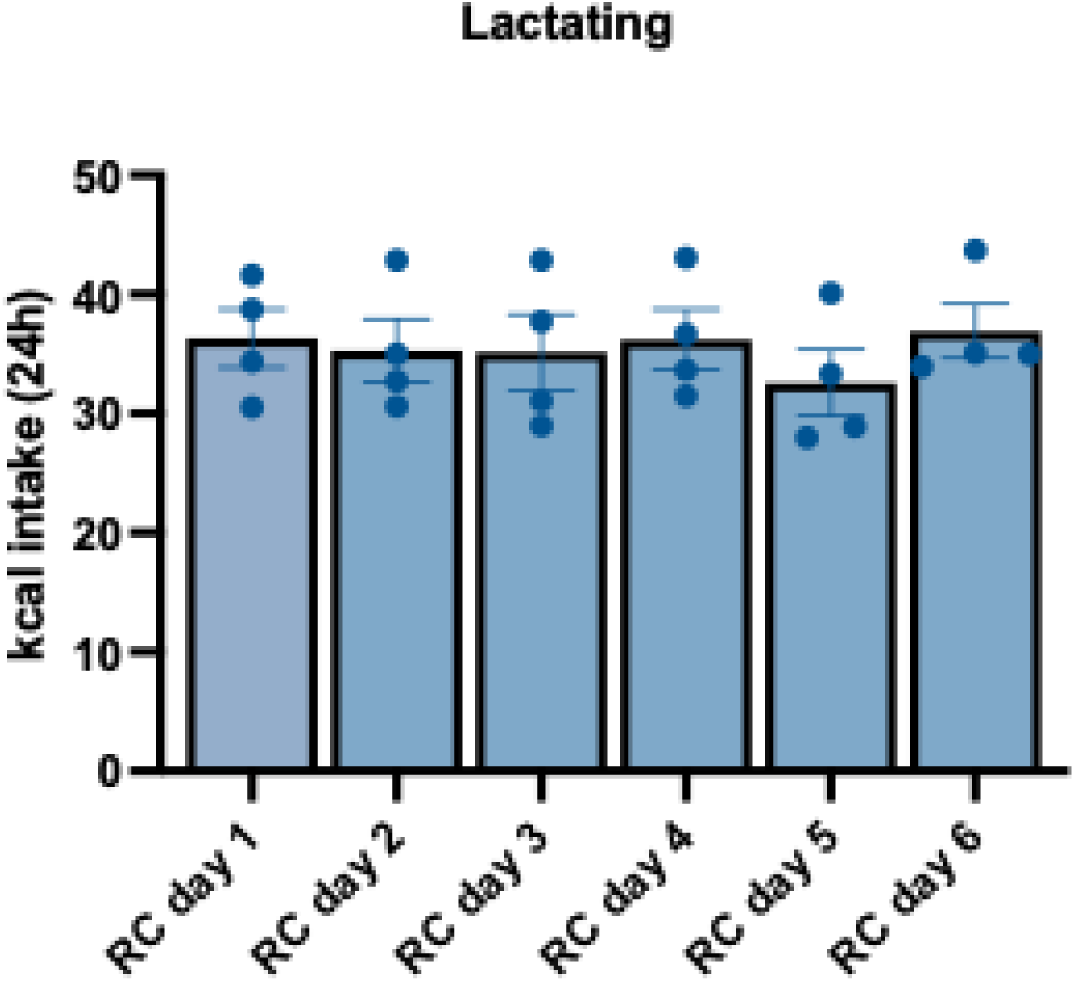
: Lactating mice consume stable amounts of calories when fed with regular chow for 6 days. Daily food intake in lactating mice provided with ad libitum access to regular chow daily.

**Extended Data Figure 2.**
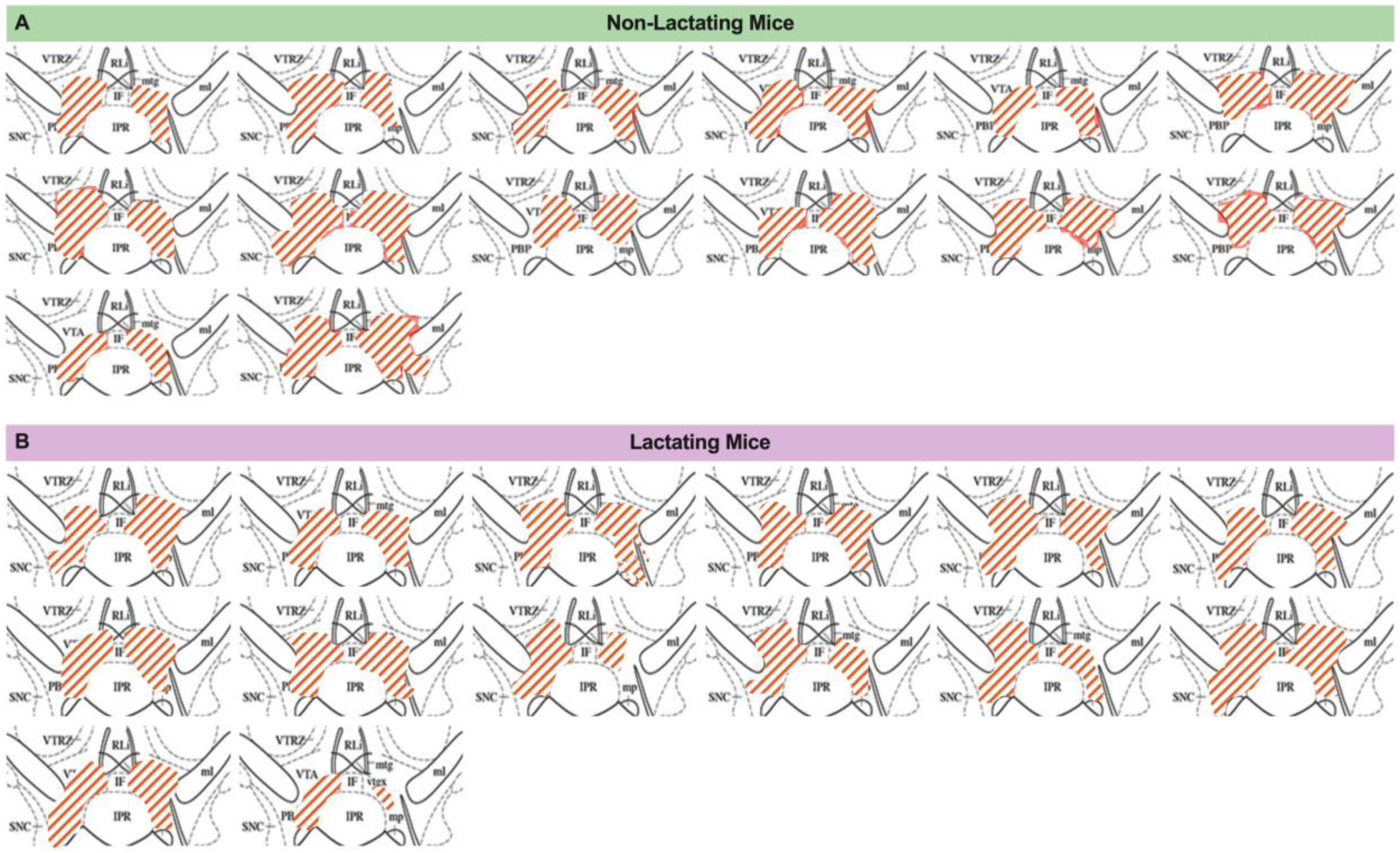
: Anatomical location of hM4Di-mCherry injections in DAT-Cre mice. (A) Schematic showing hM4D DREADD expression in VTA sections of non-lactating mice. (B) Schematic showing hM4D DREADD expression in VTA sections of lactating mice.

**Extended Data Figure 3.**
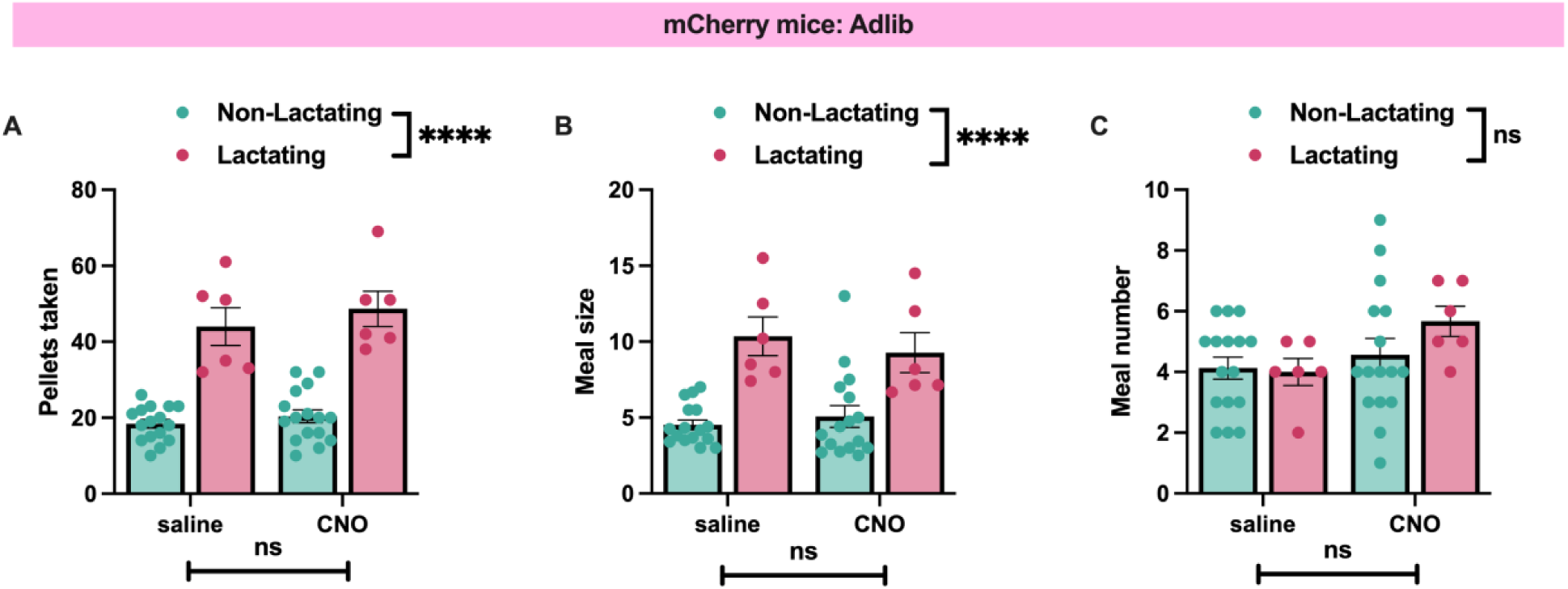
: CNO does not alter feeding behavior in non-lactating or lactating mice expressing mCherry control virus in VTA dopamine neurons. (A) Pellets consumed in non-lactating and lactation mice expressing control mCherry virus in VTA dopamine neurons following injections of saline (200ul, i.p.) or CNO (1mg/kg, i.p.; 200ul). (B) Meal size in non-lactating and lactating mice expressing control mCherry virus in VTA dopamine neurons following injections of saline or CNO. (C) Number of meals consumed following saline or CNO injections in control mCherry expressing mice that are lactating or not-lactating. Data points represent individual mice. Data analyzed by 2-way ANOVA. ns (not significant), ****p<0.001.

**Extended Data Figure 4.**
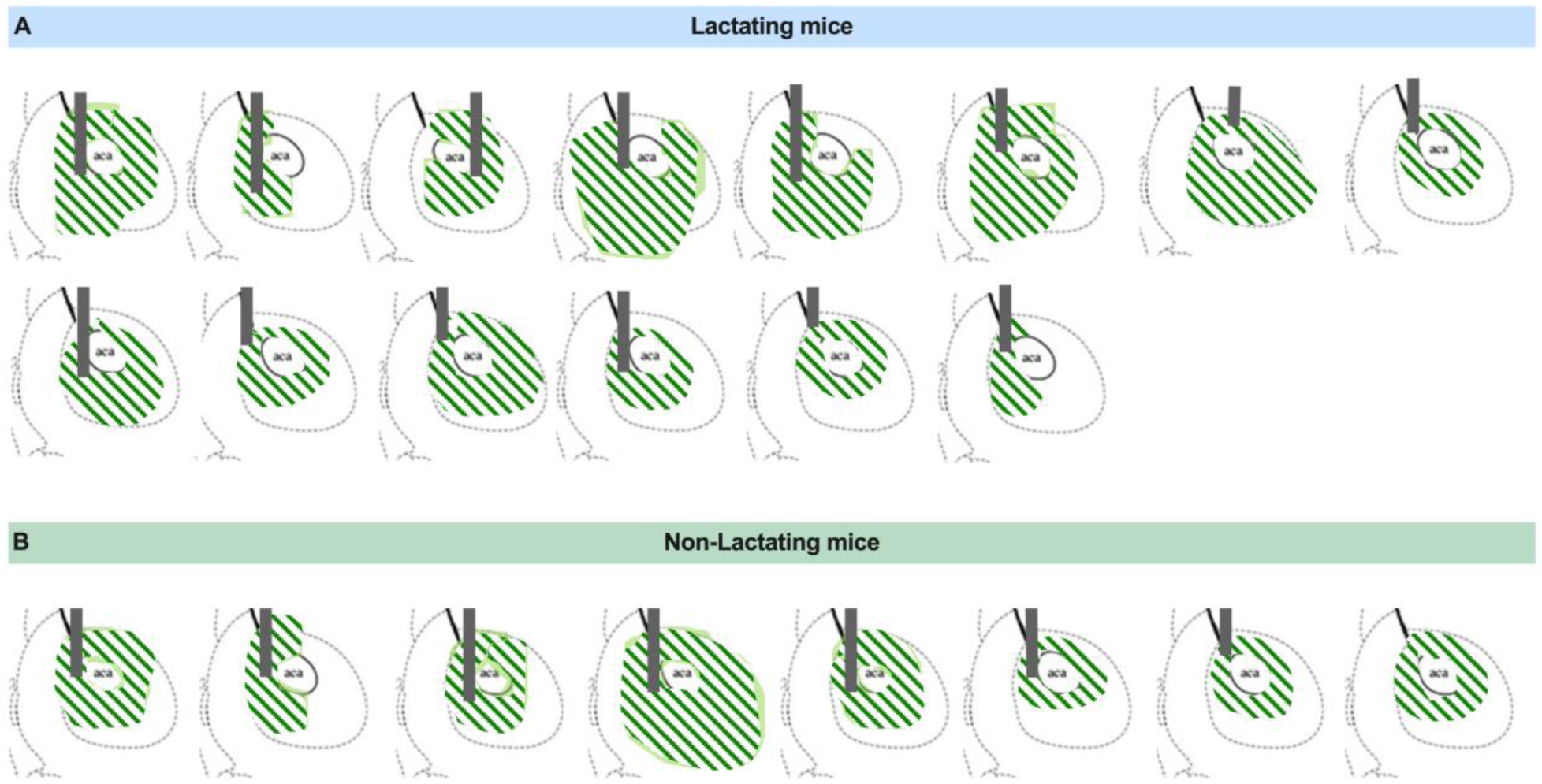
: Anatomical location of GRAB sensors and fiber optic placement in nucleus accumbens for fiber photometry experiments. (A) Schematic showing GRAB-DA expression and fiber location in striatal sections of lactating mice. (B) Schematic showing GRAB-DA expression and fiber location in striatal sections of non-lactating mice.

**Extended Data Figure 5.**
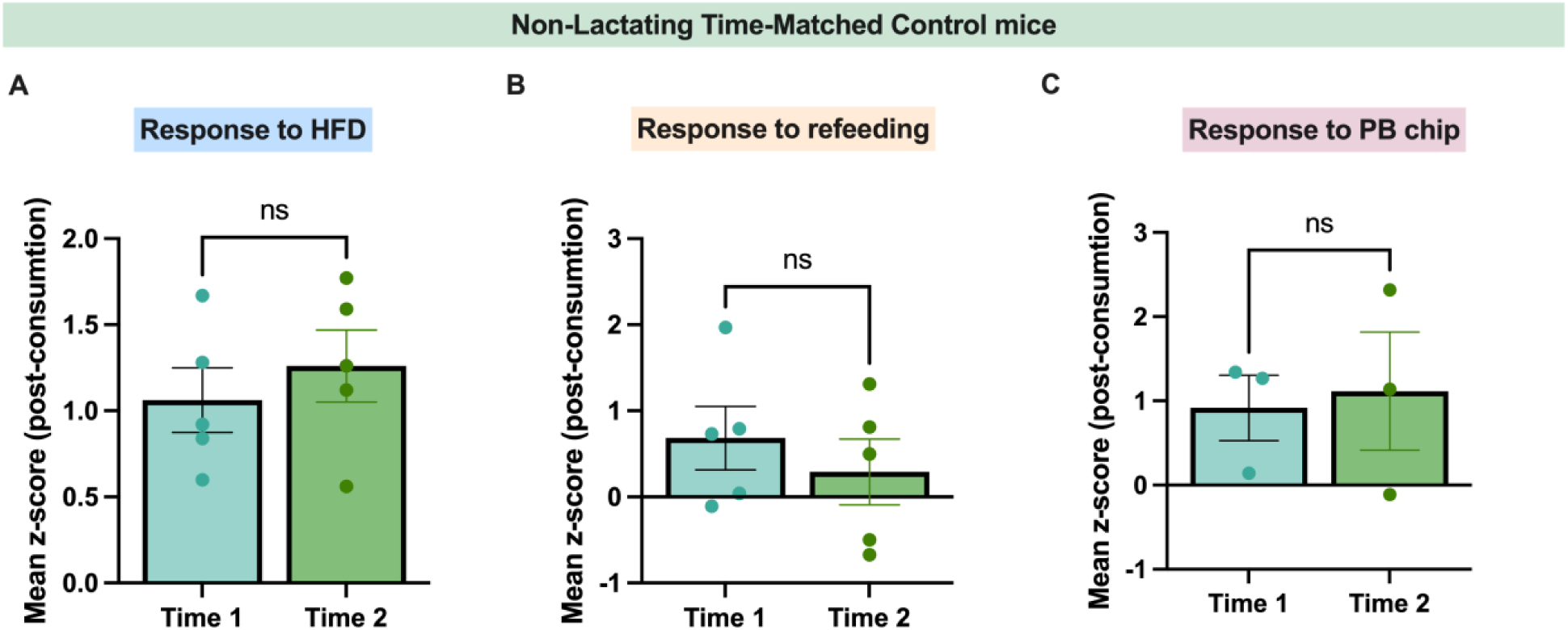
: Stable GRAB dopamine signals in time-matched control non-lactating mice. (A) Average z-score in response to high-fat consumption in non-lactating mice at two experimental time points. (B) Average z-score in response to refeeding after 10h fast in non-lactating mice at two experimental time points. (C) Average z-score in response to peanut butter chip consumption in non-lactating mice at two experimental time points. Data analyzed using student’s paired t-test. ns (not significant).

**Extended Data Figure 6.**
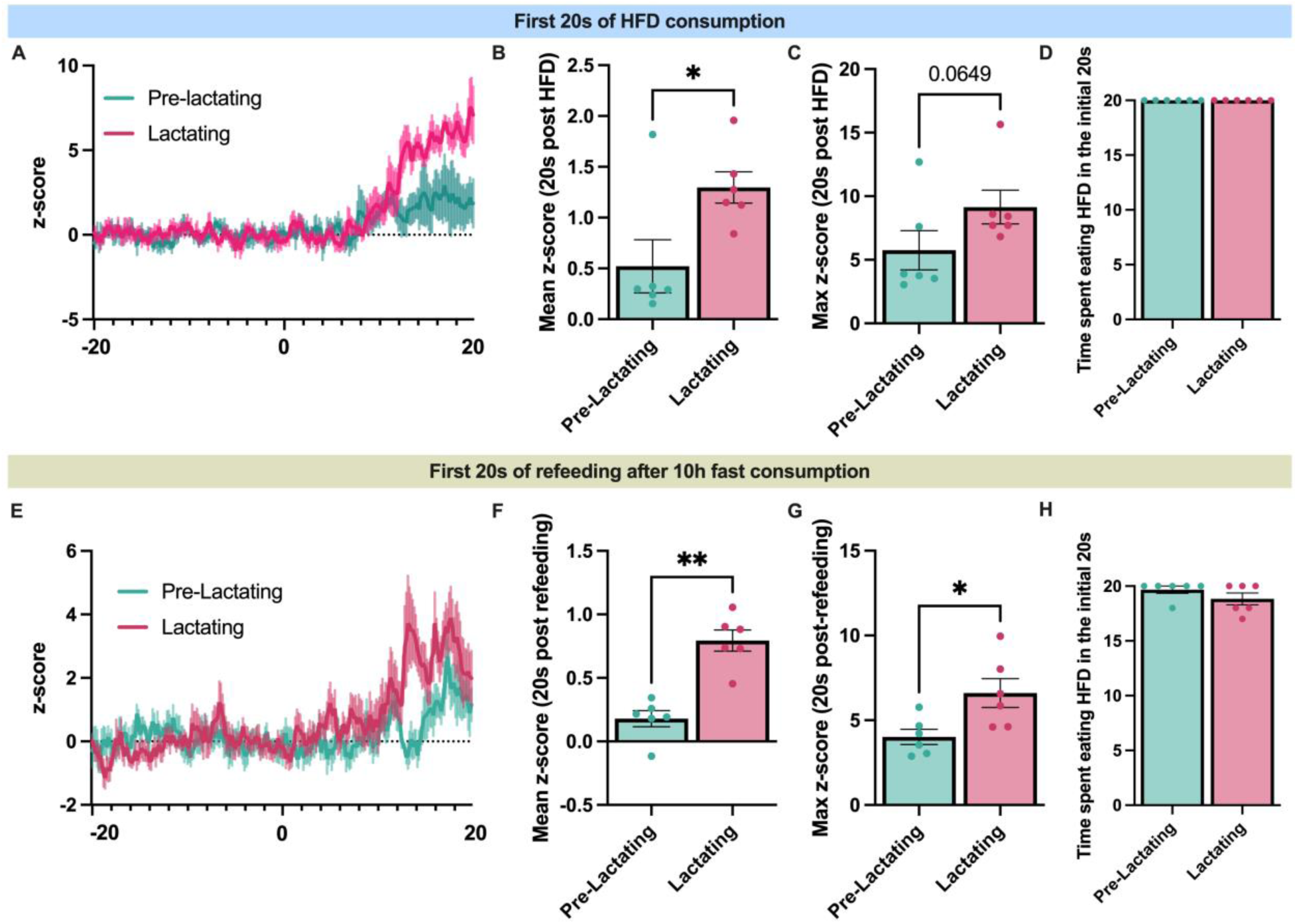
: Lactating and non-lactating mice eat for the same amount of time in the first 20 seconds of access to high fat diet or re-feeding following a ten hour fast. (A) Average trace of dopamine response following 20s consumption of high-fat diet during baseline period prior to mating and during lactation. (B) Average z-score in response to 20s of consumption of high fat diet during baseline and lactation. (C) Maximum z-score in the initial 20s of consumption of high-fat diet during baseline and lactation. (D) Total time spent eating in the initial 20s after first bite of high-fat diet in both groups, during baseline and during lactation. (E) Average trace of dopamine response following 20s of refeeding after 10h fast during baseline period prior to mating and during lactation. (F) Average z-score in response to 20s of refeeding during baseline and lactation. (G) Maximum z-score in the initial 20s of consumption of regular chow during refeeding after 10h fast. (H) Total time spent eating in the initial 20s after first bite of regular chow after 10h fast in both groups, during baseline and during lactation. Data analyzed using student’s paired t-test. *p<0.05, **p<0.01.

## Notes

### Competing Interest Statement

The authors have declared no competing interest.

